# Cyclical transcription factor AP2XII-9 is a key activator for asexual division and apicoplast inheritance in *Toxoplasma gondii* tachyzoite

**DOI:** 10.1101/2024.05.01.592006

**Authors:** Yuehong Shi, Xuan Li, Yingying Xue, Dandan Hu, Xingju Song

## Abstract

*Toxoplasma gondii* is an intracellular parasitic protozoan that poses a significant risk to pregnant women and immunocompromised individuals*. T. gondii* tachyzoites duplicate rapidly in host cells during acute infection through endodyogeny. This highly regulated division process is accompanied by complex gene regulation networks. TgAP2XII-9 is a cyclical transcription factor, but its specific role in the parasite cell cycle is not fully understood. Here, we demonstrate that TgAP2XII-9 is identified as a nuclear transcription factor and is dominantly expressed during the S/M phase of the tachyzoite cell cycle. CUT&Tag results indicate that TgAP2XII-9 targets key genes for the moving junction machinery (RON2, 4, 8) and daughter cell inner membrane complex (IMC). TgAP2XII-9 deficiency resulted in a significant downregulation of rhoptry proteins and rhoptry neck proteins, leading to a severe defect in the invasion and egress efficiency of tachyzoites. Additionally, the loss of TgAP2XII-9 correlated with a substantial downregulation of multiple IMC and apicoplast proteins, leading to disorders of daughter bud formation and apicoplast inheritance, and further contributing to the inability of cell division and intracellular proliferation. Our study reveals that TgAP2XII-9 acts as a critical S/M-phase regulator that orchestrates the endodyogeny and apicoplast division in *T. gondii* tachyzoite. This study contributes to a broader understanding of the complexity of the parasite’s cell cycle and its key regulators.

Significance: The intracellular apicoplast parasite *Toxoplasma gondii* posts great threat to the public health. The acute infection of *T. gondii* tachyzoite relies on efficient invasion by forming a moving junction structure and also fast replication by highly regulated endodyogeny. This study shows that an ApiAP2 transcription factor TgAP2XII-9 acts as an activator for the S/M-phase gene expression, including genes related to daughter buds and moving junction formation. Loss of TgAP2XII-9 results significant growth defects and disorders in endodyogeny and apicoplast inheritance of the parasites. Our results provide valuable insights into the transcriptional regulation of parasite cell cycle and invading machinery in *T. gondii*.

## INTRODUCTION

*Toxoplasma gondii* is an intracellular parasitic protozoan of the phylum Apicomplexa, which can infect humans and almost all warm-blooded animals worldwide (1). It poses a significant risk to pregnant women and immunocompromised individuals, causing severe complications and posing a public health concern worldwide (1). *T. gondii* has a complex heteroxenous life cycle comprising asexual and sexual stages. The asexual stage in the warm-blooded intermediate host consists mainly of fast-replicating tachyzoites and quiescent bradyzoites. Continuous asexual replication through endodyogeny of tachyzoites is the main cause of clinical symptoms in humans and animals (2). The sexual reproductive cycle is restricted to the intestines of the felid definitive host, during which parasites are amplified by endopolygeny to produce multiple daughter cells (3). The growth of *T. gondii* tachyzoites in cells involves a complete set of lytic cycles, including invasion, intracellular replication, and egress (2). Each tachyzoite in the host cell produces two daughter parasites via budding, a division pattern known as endodyogeny (4). Notably, the cell cycle of *T. gondii* can be categorized into G1, S, M, and C phases, lacking the classical definition of the G2 phase (5–7). During this process, the division and separation of tachyzoite organelles are allocated to the daughter cells in a highly coordinated manner, which ultimately ensures that each daughter parasite obtains a full set of organelles (8). Remarkably, the cell cycles of tachyzoites within the same parasitophorous vacuole are typically synchronized, revealing a precise and robust regulatory system to coordinate the synchronized division of these parasites (8). Transcriptomic analysis has shown differences in gene expression at different developmental stages or cell cycles (6). In *T. gondii*, there is a "just-in-time" expression pattern in which the parasite produces transcripts and proteins of certain genes through strict regulation when they are required for their function (7, 9).

Apicomplexan parasites have a unique family of transcription factors that are characterized by the possession of one or more plant-like AP2 DNA-binding domains. These apicomplexan AP2 transcription factors (ApiAP2) can bind to specific promoter sequences and regulate the expression of target genes (10). A total of 67 ApiAP2s have been identified in *T. gondii*, of which 24 are considered to be cyclically expressed ApiAP2s (9). TgAP2XI-4, TgAP2IX-4, TgAP2IX-9, TgAP2IV-3, TgAP2IV-4, and TgAP2IX-9 have been shown to be involved in the regulation of gene expression and the formation of tissue cysts (11–15). TgAP2X-4 is crucial for the growth of *T. gondii* during the acute stage of infection and is able to regulate the expression of cell cycle genes in tachyzoites (16). TgAP2X-5 has been proven to indirectly regulate the promoter of virulence genes expressed in the S/M phase, mainly in synergy with TgAP2XI-5 (7). TgAP2IX-5 was found to be a key transcriptional regulator of the asexual cell cycle and plastid division in *T. gondii* (17, 18). Knockdown of the cyclically regulated TgAP2IX-5 also results in flexibility of the cell cycle pattern from endodyogeny to endopolygeny (17). Chromatin immunoprecipitation results demonstrated that TgAP2IX-5 binds to the promoters of many of the inner membrane complex (IMC) genes and five cyclically expressed ApiAP2s (TgAP2IV-4, TgAP2III-2, TgAP2XII-9, TgAP2X-9, and TgAP2XII-2) (19). The individual gene knockout experiment showed that TgAP2III-2 is dispensable, while TgAP2IV-4, TgAP2X-9, TgAP2XII-2, TgAP2XII-9 may be essential genes (20). TgAP2IV-4 was further identified as a key suppressor of bradyzoite genes, and deletion of this gene results in the expression of a subset of bradyzoite-specific proteins during replication of tachyzoites (20). TgAP2XII-2 is associated with TgAP2IX-4 and microrchidia (MORC) and represses merozoite-specific gene expression (21, 22). TgAP2X-9 is repressed by TgAP2IX-5 (19) and may be phosphorylated by the CDK-related kinase TgCrk4 in the control of G_2_ phase (23). However, the role of TgAP2XII-9 remains unclear.

Here, we identify TgAP2XII-9 as an important S/M-phase transcription factor that activates the transcription of key genes in the moving junction complex and IMCs during daughter cell formation. Knockdown of TgAP2XII-9 leads to significant growth defects as well as disorders in endodyogeny and apicoplast inheritance. Our results provide valuable insights into the transcriptional regulation of parasite cell cycle and invading machinery in *T. gondii*.

## RESULTS

### TgAP2XII-9 expression in the nucleus during the S/M phase of the cell cycle in tachyzoites

Among a total of 67 *T. gondii* ApiAP2s, TgAP2XII-9 was recognized as a potential cell cycle-related ApiAP2 transcription factor (9, 24, 25). Amino acid sequence analysis showed that TgAP2XII-9 contains one conserved AP2 domain located at the 402-455 amino acids of its N-terminus. The AP2 domain of TgAP2XII-9 shares over 80% amino acid homology with other apicomplexans, including *Cyclospora cayetanensis* (cyc_03452), *Eimeria acervulina* (EAH_00021030), *Eimeria maxima* (EMWEY_00029950), *Sarcocystis neurona* (SN3_01800170), *Eimeria tenella* (ETH2_0411800) and *Eimeria necatrix* (ENH_00052610), and *Neospora caninum* (NCLIV_066800) (Fig. 1A and B). A previous transcriptomics study demonstrated that the expression of TgAP2XII-9 is characterized as highly dynamic during the parasite cell cycle, with peak expression during the S/M phase (9) (Fig. 1C). To confirm that TgAP2XII-9 protein is expressed during the cell cycle, we strategically fused the mAID-3HA tag to the C-terminal end of its endogenous locus, enabling tagging under their native promoters (Fig. 1D). The correct insertion of endogenous tags and single clones was confirmed by PCRs (Fig. 1E). Immunofluorescence assays (IFA) using cell cycle markers demonstrated that TgAP2XII-9 is located in the nucleus and is mostly expressed in the S/M phase (Fig. 1F and G). The expression of TgAP2XII-9 was detected during centrosome divisions rather than in individual centrosomes (identified by TgCentrin1; a marker of the outer core of the centrosome, Fig. 1G), indicating that TgAP2XII-9 is expressed in the S phase rather than the G1 phase. Moreover, the expression of TgAP2XII-9 was also detected in the M phase, as indicated by the budding marker TgIMC1 (Fig. 1F). Therefore, the expression of TgAP2XII-9 protein is rigorously controlled by the cell cycle.

**Figure 1.**
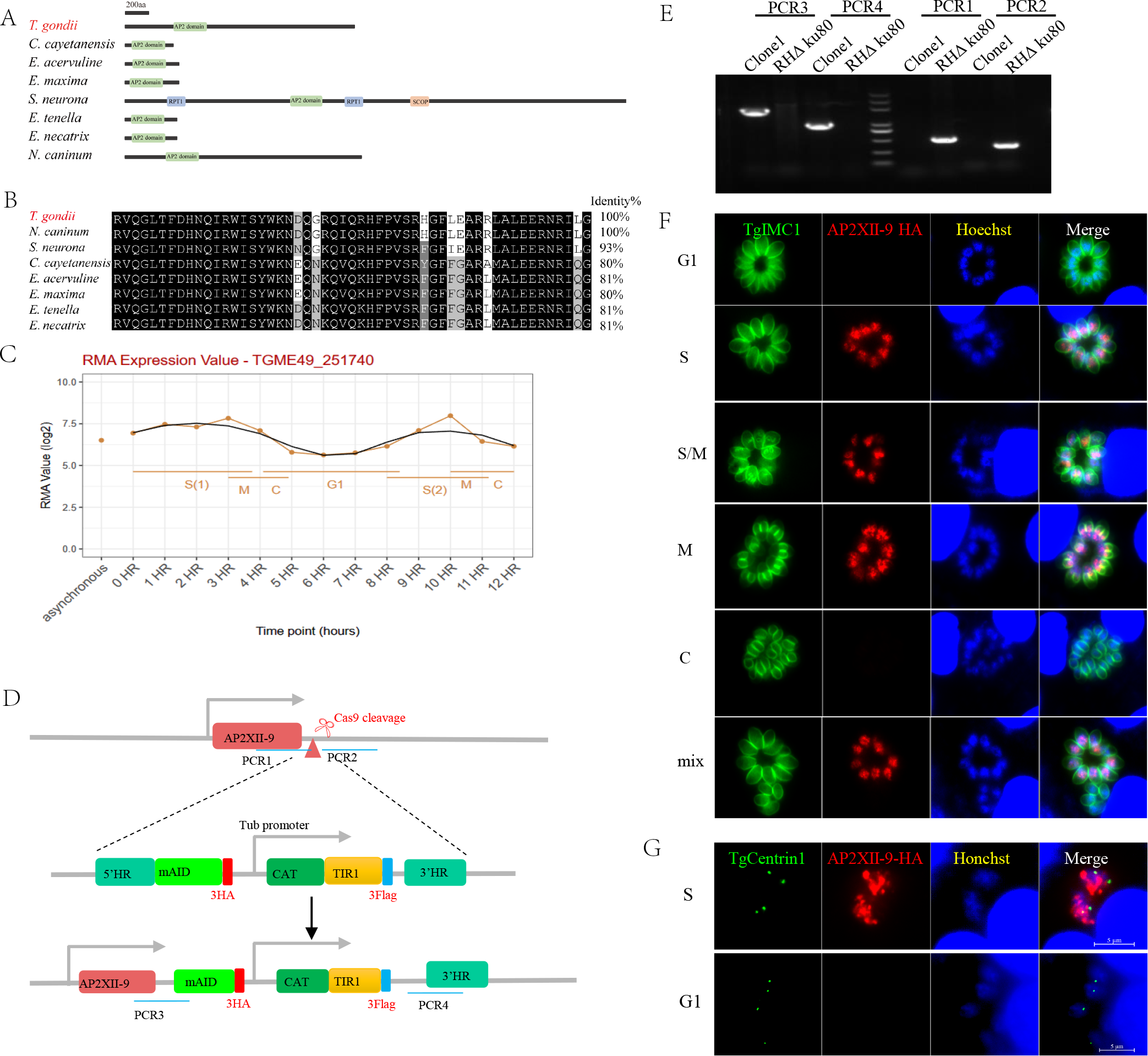
TgAP2XII-9 is a cell-cycle-regulated protein. (A) Schematic representation of the conserved AP2 domain-containing proteins from *T. gondii* (TGGT1_251740), *Neospora caninum* (NCLIV_066800), *Sarcocystis neurona* (SN3_01800170), *Cyclospora cayetanensis* (cyc_03452), *Eimeria acervulina* (EAH_00021030), *Eimeria maxima* (EMWEY_00029950), *Eimeria tenella* (ETH2_0411800) and *Eimeria necatrix* (ENH_00052610). Domains are predicted by SMART. (B) Multiple sequence alignments of the AP2 domains in apicomplexan parasites. The AP2 domain sequences from TgAP2XII-9 were aligned with their homologs. Regions of high identity and similarity between AP2 domain sequences are shown as black and gray columns, respectively. The percent homology of TgAP2XII-9 with each AP2 domain is shown at the end of the alignment. (C) *T. gondii* RH cell cycle microarray expression profiles from ToxoDB (9): X-axis time points in hours, Y-axis: RMA Normalized Values (log base 2) or expression percentile values. (D) Diagram showing the strategy for tagging TgAP2XII-9 with AID-3HA in wild parasites. The location of the primers used for the integrated PCR is indicated. (E) Confirmation of recombinant and clonal lines of iKD TgAP2XII-9 parasites by PCR. (F-G) Subcellular localization of the TgAP2XII-9 protein during the tachyzoite cell cycle. The cell cycle stage is shown on the left side of the images. Tachyzoites were co-stained using mouse anti-HA (red), rabbit anti-inner membrane complex-1 (IMC1, green) or rabbit anti-Centrin1(green) antibodies. Hoechst33258 was used to stain nuclei. IMC1 staining was used to demonstrate the tachyzoite cell cycle because the IMC1 staining pattern does not distinguish well between G1 and S phases. Division of the centrosome initiated during the S phase was identified by staining of TgCentrin1. Parasites in G1 contain single centrosomes, whereas those in S-phase are duplicated. C, cytokinesis; G1, gap phase; M, mitotic phase; S, synthesis phase.

### TgAP2XII-9’s critical role in tachyzoite growth and intracellular replication

To assess the functions of TgAP2XII-9 in *T. gondii*, an indole-3-acetic acid (IAA) degradation (mAID) system was used to generate an inducible knockdown (iKD) of TgAP2XII-9 parasites (iKD TgAP2XII-9) (Fig. 1D, E). Immunofluorescence assays confirmed that the addition of IAA for 3 h resulted in the degradation of TgAP2XII-9 protein in tachyzoites (Fig. 2A). To comprehensively evaluate the viability of TgAP2XII-9-deficient parasites, we monitored the formation of plaques during a continuously maintained 7-day culture. The results showed that the parasites were unable to form plaques in host cells upon IAA treatment, whereas untreated parasites formed noticeable plaques (Fig. 2B and C). The growth of *T. gondii* tachyzoites in cells involves a complete set of lytic cycles, including invasion, intracellular replication, and egress. The reduction in plaque formation may be caused by impairment of one or more steps of the lytic cycle. Thus, we next sought to investigate the role of TgAP2XII-9 in the lytic cycle biology of *T. gondii*. The invasion process of the parasites was first assessed, and the invasive ability of iKD TgAP2XII-9 parasites treated with IAA was significantly reduced compared to untreated parasites (7%, *P* <0.01) (Fig. 2D). Subsequently, the calcium ionophore A23187 was used to assess the egress ability of the parasites, and the results showed a significantly decreased egress ratio (33.3%, *P* <0.05) in the iKD TgAP2XII-9 parasites after 30 h of IAA treatment (Fig. 2E). For intracellular replication, degradation of TgAP2XII-9 led to a severe decrease in the intracellular replication capacity of parasites having 1 or 2 tachyzoites per vacuole, whereas parasites without IAA treatment mostly had 4 and 8 tachyzoites per vacuole (Fig. 2F). It is worth noting that the replication defects of the parasites caused by degradation of the TgAP2XII-9 protein are not irreversible. After the addition of IAA treatment for 16 h to induce degradation of the TgAP2XII-9 protein, the parasites were placed in a culture medium without IAA for another 0, 3, 6, and 20 h, respectively. The results showed that the proliferation ability of parasites treated in this way significantly recovered during this period (Fig. 2G). Interestingly, about 8.7% of the parasitic vacuoles (PVs) exhibited significant asynchronous division after removal of IAA (*P* <0.001, Fig. 2H-I, J), implying inconsistent growth cycles of parasites within the same PV. These results indicate that TgAP2XII-9 expression is crucial for the lytic cycle of tachyzoites.

**Figure 2.**
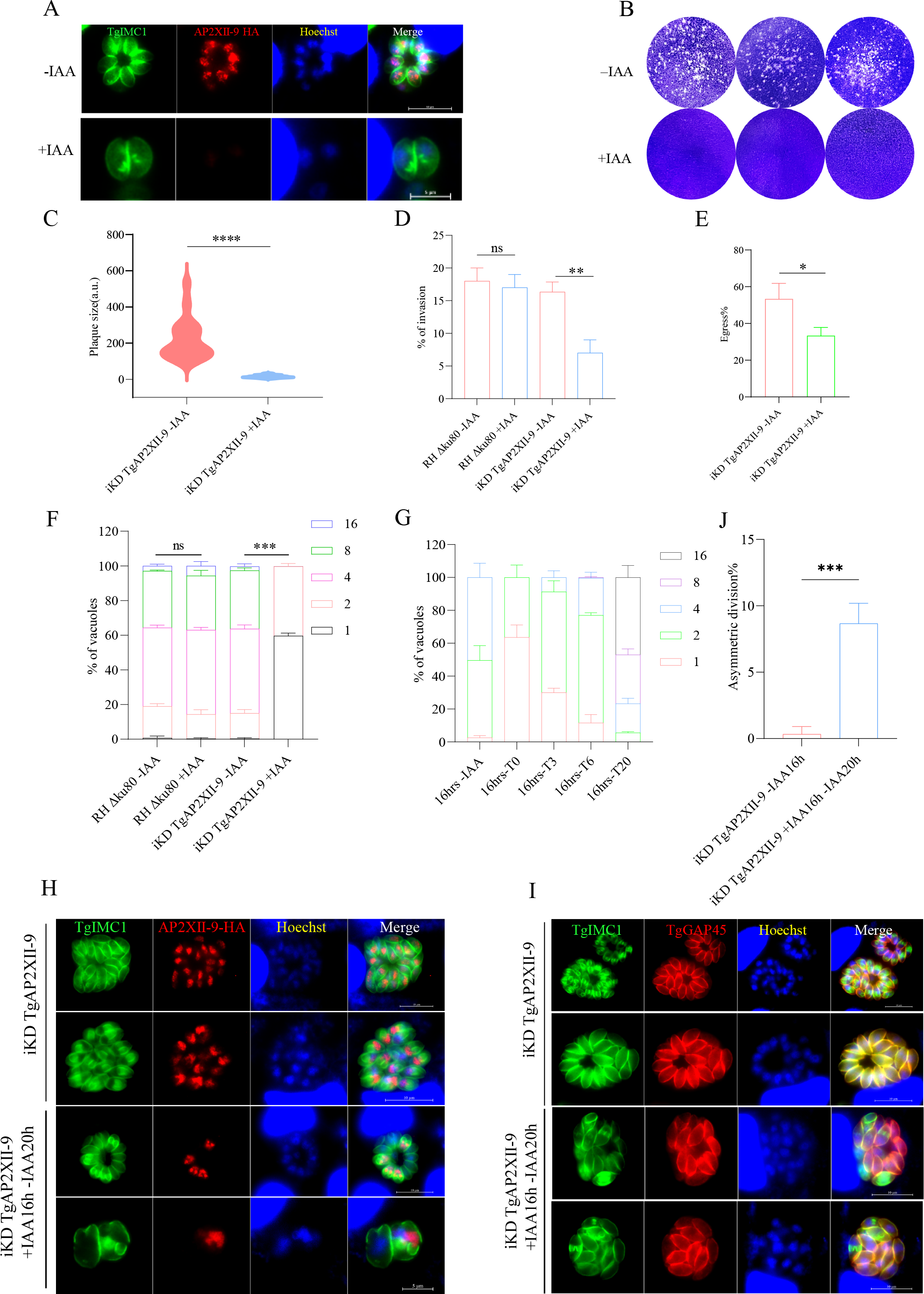
TgAP2XII-9 is essential for the replication of *T. gondii* tachyzoites. (A) Fluorescence microscopy of intracellular iKD TgAP2XII-9 parasites after treatment with 500 μM IAA or vehicle. Tachyzoites were co-stained with mouse anti-HA (red) and rabbit anti-TgIMC1 (green) antibodies. Nuclei were stained with Hoechst (blue). Scale bars = 5 μm. (B-C) Plaque assay of iKD TgAP2XII-9 parasites grown on IAA- or vehicle-treated HFF cells for 7 days. Plaque areas were measured and counted by Photoshop C6S software (Adobe, USA). (D) Inoculation of 1 × 10^5^ parasites onto HFF cells in 12-well plates and cultured for 24 h. IFA was performed using anti-TgGAP45 antibodies and Hoechst. The invasion ratio of TgAP2XII-9 knockdown parasites was grounded on the number of parasite-infecting cells divided by the total number of cells in one horizon. Asterisks indicate statistically significant results (p < 0.01 as determined by t-test). Data are the mean ± SD (error bars) of three independent experiments. (E) Efficiencies of A23187-induced egress of iKD TgAP2XII-9 parasites that were pretreated with or without IAA for 30 h. The average number of ruptured PVs were determined by randomly counting 100 vacuoles per slide. Means ± SD of three independent experiments were graphed. (F) Intracellular parasite replication after incubation with IAA or vector at 24 hours post-infection. Data are expressed as the mean ± SEM of three independent assays, each counting 100 total PVs per strain. (G) Assessment of intracellular growth of iKD TgAP2XII-9 parasites after 16 h of culture in Auxin-added medium, followed by 0, 3, 6, and 20 h of culture in standard medium (16hrs-T0/T3/T6/T20). Parasite nuclei were stained with hoechst33258 and the parasite inner membrane complex (IMC) was stained with TgIMC1 antibody. The number of tachyzoites in the PV was counted by three independent assays of 100 PV each. (H-I) Immunofluorescence assay of iKD TgAP2XII-9 parasites treated with auxin for 16 h and recovery of growth after auxin washout. Parasites were fixed and stained with rabbit anti-IMC1 antibody (green), mouse anti-HA antibody (red) or rabbit anti-GAP45 antibody (red). (J) The abnormal proportion of asynchronously dividing parasitic vacuoles (PVs). *: *P* < 0.05, **: *P* < 0.01, ***: *P* < 0.001, ****: *P* < 0.0001, n.s: no significant difference.

### TgAP2XII-9 regulates transcription of periodic genes and targets ROP and IMC genes

Potentially regulated genes of the transcription factor TgAP2XII-9 were investigated by RNA-seq using TgAP2XII-9 iKD parasites. A total of 1765 differentially expressed genes (DEGs) were identified after TgAP2XII-9 knockdown, including 981 downregulated genes (more than 2-fold decrease in the transcript level; *P* <0.05) and 784 upregulated genes (more than 2-fold decrease in the transcript level; *P* <0.05) (Fig. 3A; Table S2). The cell cycle expression profile of downregulated genes was analyzed using previously published data (9) and represented as a heat map (Fig. 3B). The results showed that most of the downregulated genes exhibited features of cell cycle regulation. Subsequently, we clustered these downregulated genes with similar expression trends and found that the expression profiles of these genes could be divided into three clusters (Fig. 3B, C; Table S2). Interestingly, the genes in cluster 3 showed the highest expression in S and M phases and the lowest expression in the G1 phase, which is consistent with the expression pattern of TgAP2XII-9. On the contrary, most of the genes in cluster 1 showed highest expression in the G1 phase and lowest expression in the S and M phases. Most genes in cluster 2 showed peak expression in late G1 and early S phases, whereas the low expression peaks corresponded to M and early G1 phases (Fig. 3C).

**Figure 3.**
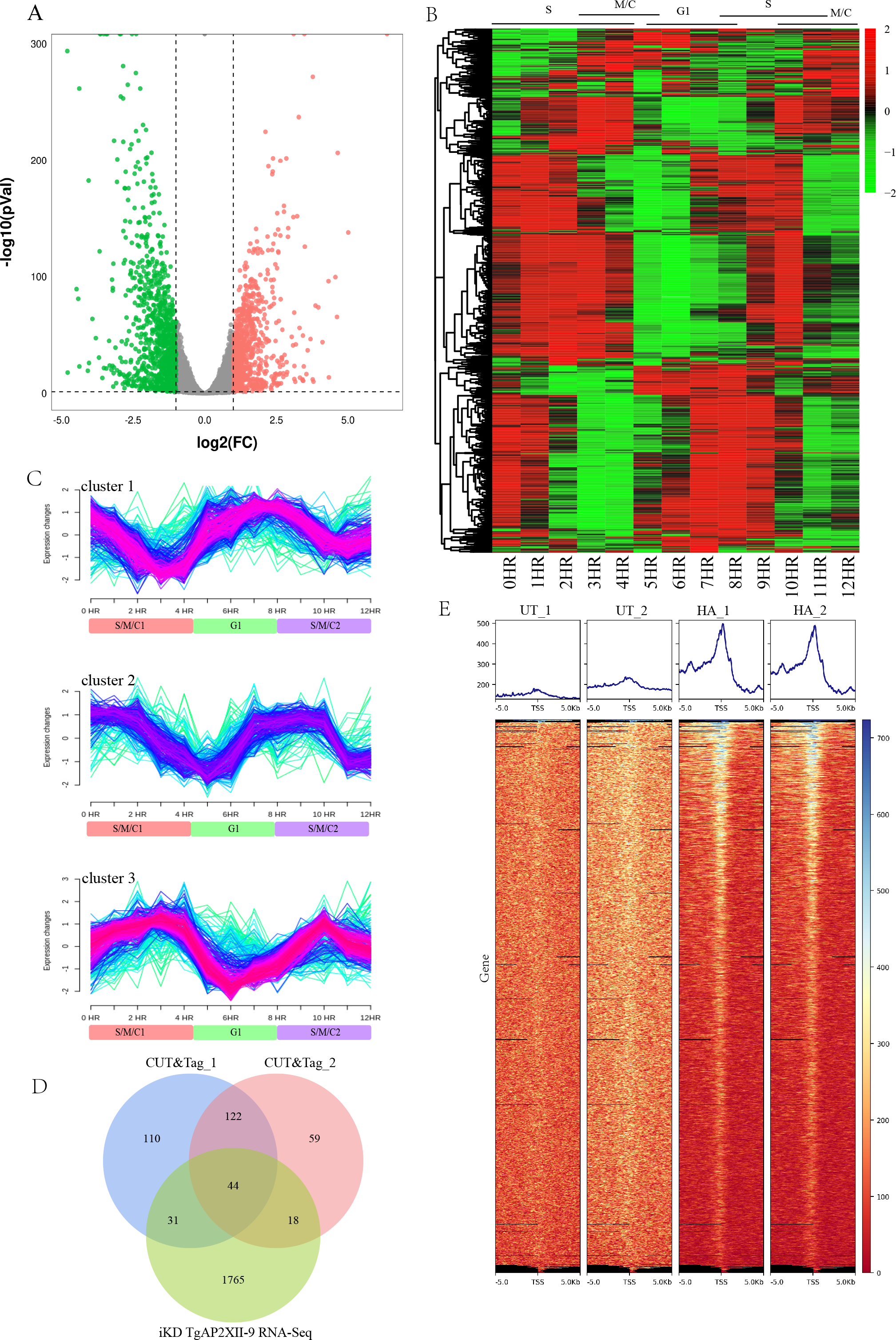
**TgAP2XII-9 controls multiple cell-cycle-regulated genes. (**A) Volcano plots showing differentially expressed genes after degradation of TgAP2XII-9 protein by IAA treatment for 24 h (n= 8,920). Green and blue dots indicate significantly downregulated and upregulated genes, respectively. (B) Heatmap showing the cell cycle expression of all individual transcripts downregulated in the iKD TgAP2XII-9 strain in the presence of 24 h of auxin treatment. Cell cycle phases (G1-S-M-C) are shown at the top. (C) Mfuzz cluster analysis illustrating changes in the expression of down-regulated genes during cell cycle progression (9). Fuzzy c-means clustering identified three distinct temporal patterns of protein expression. The x-axis represents time points in hours, while the y-axis represents the log2-transformed, normalized intensity ratios in each stage. cluster1: G1-S high expression peaks; cluster2: S-M low expression peaks. Cluster3: G1 low expression peaks. (D) Venn diagram of TgAP2XII-9-dependent DEGs intersected with the TgAP2XII-9 CUT&Tag genes. (E) Profile and heat maps of the averaged sum showing the CUT&Tag called the peaks of APXII-9 (HA) around the TSS of the parasite genes. The top panels show the average signal profile of the genomic loci centered on the TSS (±5 kb). The lower panels show heat maps of peak density around the same genomic loci. The color scale used to interpret the signal intensity is located on the right side of each graph.

To further characterize the direct target genes of TgAP2XII-9, CUT&Tag experiments were performed using the endogenously 3xHA tagged parasite and RH strain (Fig. 3D). Promoters of 251 and 320 genes were enriched for the two individual CUT&Tag experiments, respectively (Fig. 3E; Table S2). Of these, 164 genes appeared in both replicates, with a total of 44 genes were differently expressed after TgAP2XII-9 knockout. Of these 44 potential targets, eleven proteins were rhoptry proteins (ROP1, ROP4, ROP15, ROP17, ROP39, ROP47, ROP48) or rhoptry neck proteins (RON2, RON3, RON4, RON8); 14 IMCs or glideosome-associated proteins, including IMC3, IMC4, IMC10, IMC16, IMC31, IMC34, AC2, AC12, AC13, GAP40, GAP50, GAPM3,

GAPM1A, GAPM2A, and IMC localizing protein (ILP1). Moreover, all these ROPs/RONs and IMC related genes were downregulated upon TgAP2XII-9 depletion, indicating that TgAP2XII-9 binds directly to the promoters of these genes and functions as an activator.

### TgAP2XII-9 influences the invasion and egress efficiency of tachyzoites by regulating the expression of ROP genes

As TgAP2XII-9 knockout results in differential transcription of multiple periodic genes, these genes can be categorized into three major clusters. Our attention was first focused on the genes in cluster 3 because its expression pattern is similar to that of TgAP2XII-9, both of which are highly expressed in the S/M phase and downregulated in the G1 phase. GO enrichment analysis revealed that cluster 3 genes were mainly distributed in the inner member pellicle complex, pellicle, and apical part of cell (Fig. 4A). We further conducted a detailed analysis of these genes and found that most of the proteins clustered in the apical part of the cell were ROP and RON proteins (*P* < 3.26E-15). These ROP and RON proteins have a very similar periodic expression pattern to TgAP2XII-9 during the cell cycle, peaking at S and M phases and reaching a minimum at the G1 phase (Fig. 4B, C).

**Figure 4.**
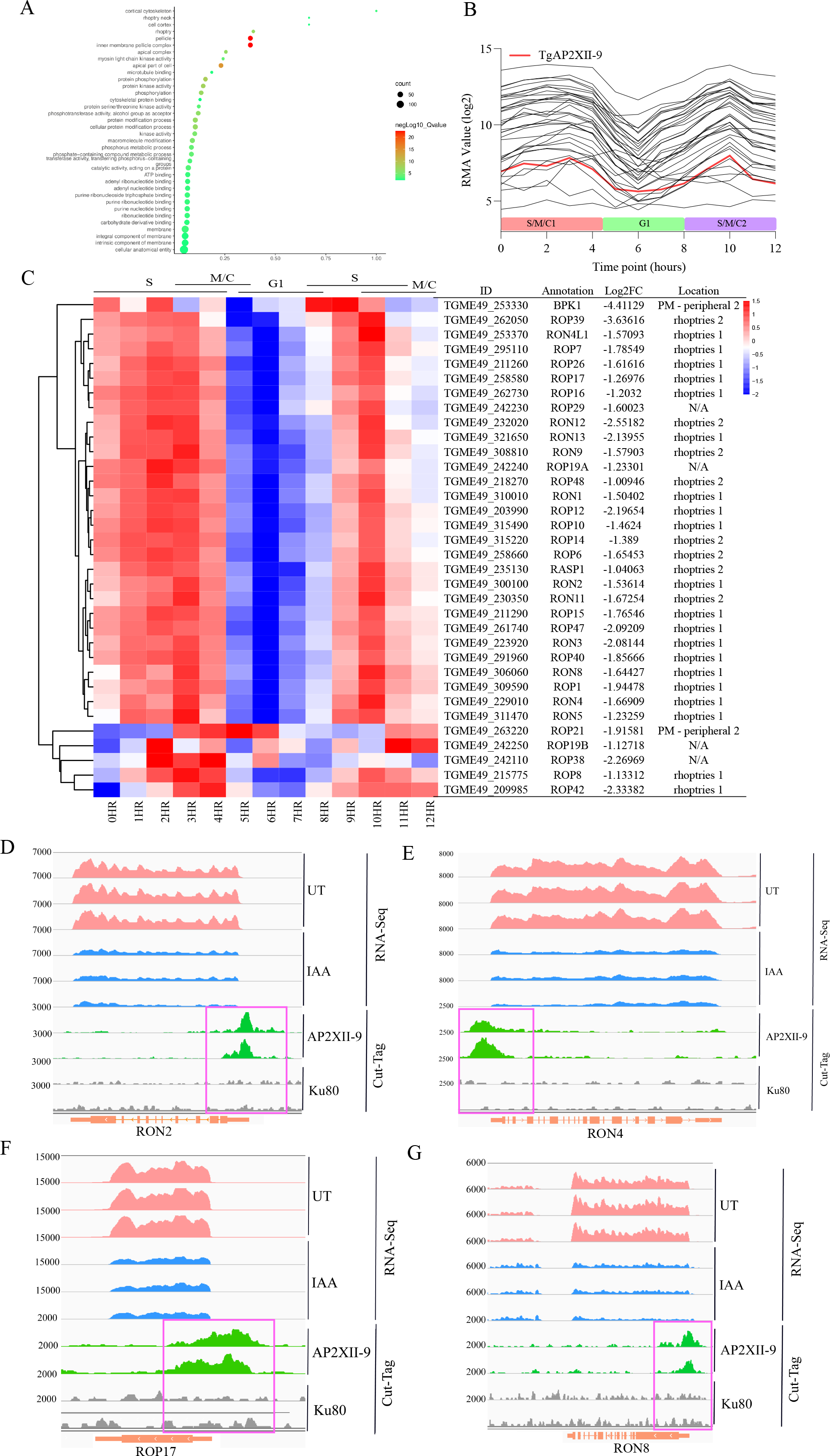
TgAP2XII-9 is required for efficient invasion of host cells. (A) Performance of functional enrichment for cluster 3 proteins by gene ontology (GO) analysis using ToxoDB annotations. Significant GO terms are shown (Benjamini < 0.1). (B) The mRNA profiles of the downregulated rhoptries (ROP) and rhoptry neck (RON) genes in AP2XII-9-depleted parasites according to the cell-cycle expression profiles (9) (source of data: ToxoDB.org); RMA, robust multiarray average; G1, G1 phase; S, synthesis phase; M, mitosis phase; C, cytokinesis phase. (C) Hierarchical clustered transcriptional heatmap showing the transcription level of downregulated ROP and RON genes (n=35) after AP2XII-9 depletion at different cell cycle phases. The top of the heatmap shows the expression scale of cell cycle phases (S-M-C-G1). IGV screenshots of the genomic regions of RON2 (D), RON4 (E), ROP17 (F) and RON8 (F). CUT&Tag profiles were obtained using antibodies directed against HA (TgAP2XII-9 tagged) in chromatin sampled from the AP2XII-9-mAID-HA strain and the RH Δku80 strain. RNA-seq data from iKD TgAP2XII-9 parasites after 24 h of treatment or no treatment with IAA are shown in blue and pink peaks. The normalized RPKM for Cut-Tag and RNA-seq reads are shown on the y-axis.

The moving junction (MJ), which consists of apical membrane antigens and RONs protein complex, is a key structure for parasite invasion into host cells (26). The deletion of TgAP2XII-9 resulted in a significant downregulation of 11 RONs. As a result, the transcription levels of all moving junction-related RONs (RON2, RON4, RON4L1, RON5, and RON8) were significantly downregulated. In addition, the promoters of ROP17, RON2, RON4 and RON8 were directly bound by TgAP2XII-9 as shown in the CUT&Tag results (Fig. 4D-G). The formation of MJ in the tachyzoite is characterized by the interaction of AMA1 (secreted by micronemes and relocalized to the parasite surface prior to invasion) and the export of RON2, with further recruitment of other RONs (27). The depletion of these RONs would greatly impair the invasion ability, and we did observe a significant reduction in the invasion rate after TgAP2XII-9 depletion. Interestingly, the bradyzoite MJ components AMA2 (Log2Fold Change = -1.41, FDR = 6.48E-22) and AMA4 (Log2Fold Change = -4.37, FDR = 2.25E-262), which are key proteins for cysts burden during the onset of chronic infection (28), were also down-regulated due to the absence of TgAP2XII-9. This suggests that the invasion ability of bradyzoites may also be affected by TgAP2XII-9, but further validation is needed.

We also identified 23 rhoptry proteins that are down-regulated upon deletion of TgAP2XII-9, and 7 ROPs whose promoters are directly bound by TgAP2XII-9, including ROP1, ROP4, ROP15, ROP17, ROP39, ROP47, and ROP48. A subset of these proteins has been fully characterized as virulence factors interacting with host cells. ROP1 was the first rhoptry protein to be identified and is essential for parasite virulence in vitro and in vivo, as well as crucial for counteracting interferon gamma-mediated innate immune restriction (29). ROP1 also binds to host Complement Component 1q Binding Protein (C1QBP), which is a regulator of autophagy and innate immunity (29). ROP17, an active rhoptry kinase localized on the external surface of the PVM, forms complexes with the key virulent factors ROP5 and ROP18. ROP17 phosphorylates host cell IRGs (immunity-related GTPases recruited to the PVM and leading to its disruption and parasite death (30)) and inhibits the polymerization of IRGs as well as contributing to the virulence of *T. gondii* (31). ROP39 is another *T. gondii* virulence factor associated with ROP5B, which directly targets host Irgb10 and inhibits homodimer formation of GTPase by reducing IRG proteins loading onto the PVM (32). ROP47 is secreted from rhoptry bulb and relocalizes to the host cell nucleus, and may play a role in manipulating host cell gene transcription (33). However, deletion of ROP47 or ROP48 in a type II strain did not show a major influence on in vitro growth or virulence in mice (33). Generally, these results demonstrate that TgAP2XII-9 controls the expression of core genes for MJ and parasite virulence factors, which suggests that TgAP2XII-9 contributes to parasite invasion and virulence.

### TgAP2XII-9 controls the expression of multiple IMC genes and is crucial for the formation of daughter buds

In addition to the ROP protein, a large number of cluster 3 genes were also enriched in the inner member pellicle complex and pellicle (Fig. 4A), and the majority of these genes were found to be genes encoding IMC proteins of tachyzoites (*P* < 2.15E-23). The IMC is an important organelle in *T. gondii* that plays a key role in parasite motility, invasion, egress, and replication (2, 34) . IMC can be classified as proteins that are specifically localized to the maternal IMC, the daughter bud IMC, or both (35).

Depletion of the TgAP2XII-9 protein resulted in the downregulation of 52 IMC proteins (more than a 2-fold decrease in the transcriptional level; P<0.05), exhibiting a cyclical expression pattern consistent with that of TgAP2XII-9, with peak expression in S and M phases and reaching the bottom in the G1 phase (Fig. 5A, B). Among these downregulated IMC proteins, more than 10 proteins are known to be localized in daughter tachyzoites, including IMC1, IMC3, IMC6, IMC29, IMC30, IMC31, IMC34, IMC35, IMC36, IMC44, AC12, AC13, ISP1 and ISP2 (Fig. 5B) (35–39). Of these, 7 proteins (IMC29, IMC30, IMC31, IMC34, IMC35, IMC36 and IMC44) are restricted to the IMC body of daughter buds, and 2 proteins (AC12 and AC13) are localized only to the apical cap of daughter buds (35). IMC29 has been identified as an important early daughter bud component for replication (35). CUT&Tag results also showed that the promoters of daughter cell-restricted IMC31, IMC34, AC12, and AC13 bind directly to TgAP2XII-9 (Fig. 5C-F).

**Figure 5.**
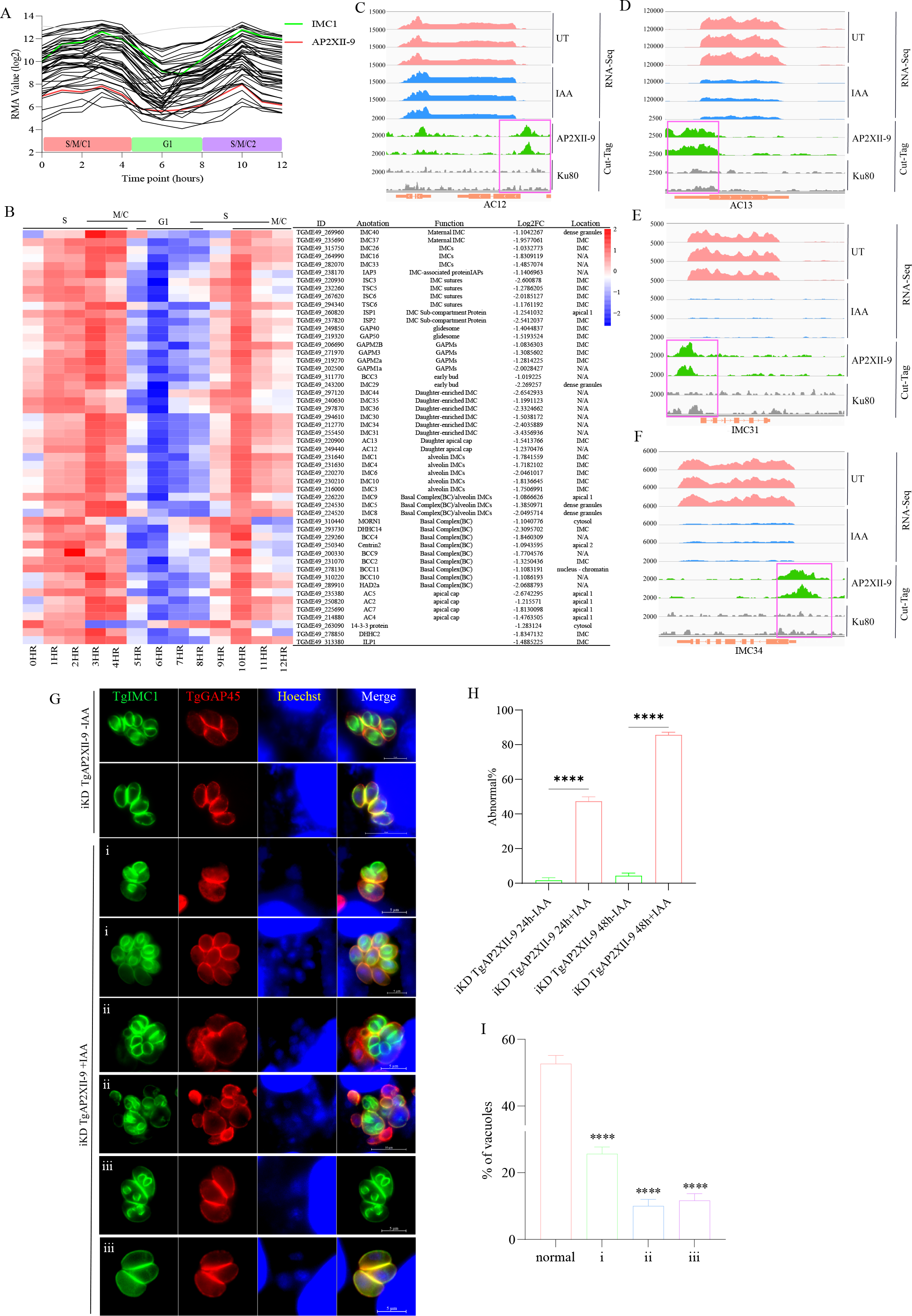
TgAP2XII-9 controls the expression of key genes involved in daughter parasite formation. (A) The mRNA expression showing downregulation of IMC genes after TgAP2XII-9 depletion (Log2 FC <-1; P-value < 0.01). Their transcript was abundant in various cell cycles of parasites, namely S, M, C, and G1 phases; RMA, robust multiarray average; G1, G1 phase; S, synthesis phase; M, mitosis phase; C, cytokinesis phase. (B) Heat map of the cell cycle expression profile for 52 IMC-associated genes transcripts downregulated (Log2 FC <-1; P-value < 0.01) in TgAP2XII-9-deficient parasites. Scale of the cell cycle phase is shown at the top of the heat map (S-M-C-G1). IGV screenshots of the genomic region of AC12 (C), AC13 (D), IMC31 (E), and IMC34 (F). CUT&Tag profiles were obtained using antibodies directed against HA (AP2XII-9-tagged) in chromatin samples of the iKD TgAP2XII-9 strain and the RH Δku80 strain. RNA-seq data from iKD TgAP2XII-9 strain parasites after 24 h of treatment or no treatment with IAA are shown in blue and pink peaks. The normalized RPKM for Cut-Tag and RNA-seq reads are shown on the y-axis. (G-H) IFA showing the patterns of daughter cell budding (stained with TgIMC1) in iKD TgAP2XII-9 parasites treated with or without IAA for 24 or 48 h. Different patterns of abnormal budding were observed: i) non-synchronous division, ii) abnormal number of daughter budding, and iii) the appearance of an odd number of tachyzoites in a PV. Nuclei were stained with Hoechst (blue). The scale bar is shown on the lower right side of each image. (I) Statistical analysis of the percentage of PVs in each type of abnormal divisions (types i, ii and iii) compared to the total abnormal PVs after 24 h of treatment with IAA. Means ± SD of three independent experiments, \**P*<0.05, \*\**P*<0.01, \*\*\**P*<0.001, Student’s *t*-test. (J) Statistical analysis of the percentage of abnormally divided PVs after 24 and 48 h of treatment with IAA.

Considering that the depletion of TgAP2XII-9 resulted in a significant downregulation of a large number of IMC proteins related to the formation of daughter buds in *T. gondii,* we further used IMC1 as a marker to verify whether the division of the TgAP2XII-9 deficient parasites was affected. After 24 and 48 h of treatment with IAA, the iKD TgAP2XII-9 parasites exhibited disordered daughter divisions (Fig. 5G and H). Three different types of unnatural divisions were mainly observed: i) non-synchronous division characterized by different budding states (budding, no budding, just budding, and complete daughter cell formation) of tachyzoites in the same PV; ii) endopolygeny with more than 2 daughter buds within a parasite (>2 daughter buds per maternal parasite); and iii) the presence of an odd number of tachyzoites in a PV (Fig. 5G). Among them, the proportion of abnormal divisions was 26%, 10% and 12% in cases i, ii and iii, respectively, after 24 h of treatment with IAA (Fig. 5I). When the IAA treatment was extended to 48 h, the parasites were observed to grow continuously and these defects seemed to accumulate over time. Quantification showed 86% of abnormal vacuoles after 48 h of IAA treatment (Fig. 5H). These results indicate that TgAP2XII-9 significantly impairs the division of tachyzoites by regulating the transcription of various budding-related IMC genes, which ultimately leads to limited intracellular replication in parasites.

### TgAP2XII-9 regulates apicoplast gene expression and its division

According to the cyclical expression profile, the genes downregulated in TgAP2XII-9 deletion parasites were categorized into three clusters (cluster 1-3) (Fig. 3C). Previous GO analysis of the cluster 3 gene showed enrichment in IMC, ROP, and RON proteins (Fig. 4A). The downregulation of these genes may be responsible for abnormal division, intracellular replication stagnation, and reduced invasion and egress efficiency in the parasites. Considering that the expression peaks of genes from cluster 1 to cluster 3 progressed over time, we speculate that TgAP2XII-9 may directly or indirectly regulate gene expression in clusters 1-3.

Therefore, we further performed GO analysis on the cluster1 genes and found that a total of 18 apicoplast genes were enriched (Fig. 6A; p-value< 2.11E-11). These genes showed the highest expression in the G1 phase (5-9 h) and the lowest expression in the S and M phases (Fig. 6B and C). Most of these apicoplast genes (9/18) are key genes in the type II fatty acid synthesis (FASII) pathway, including acetyl-CoA carboxylase (ACC1), enoyl-acyl carrier reductase (ENR), apicoplast triosephosphate translocator (APT1), acyl carrier protein (ACP), glycerol 3-phosphate acyltransferase (ATS1), and 4 pyruvate dehydrogenase complex subunits (PDH-E2, PDH-E1α, PDH-E1β and PDH-E3I) (Fig. 6B and C) (40, 41). The FASII pathway is a de novo synthesis pathway for fatty acids (FA) located in the apicoplast of *T. gondii*, and various components in this pathway are crucial for apicoplast formation and parasite replication (19, 40–42). Since the deletion of TgAP2XII-9 leads to the downregulation of many apicoplast genes, particularly those in the FASII pathway, a further desire was made to verify whether apicoplast division would be affected. By using IFAs conducted with anti-*T. gondii* ACP and ENR (apicoplast markers) antibodies, we found that the iKD TgAP2XII-9 parasite’s apicoplast exhibited genetic disorders after 24 h of treatment with IAA (Fig. 6D).

**Figure 6.**
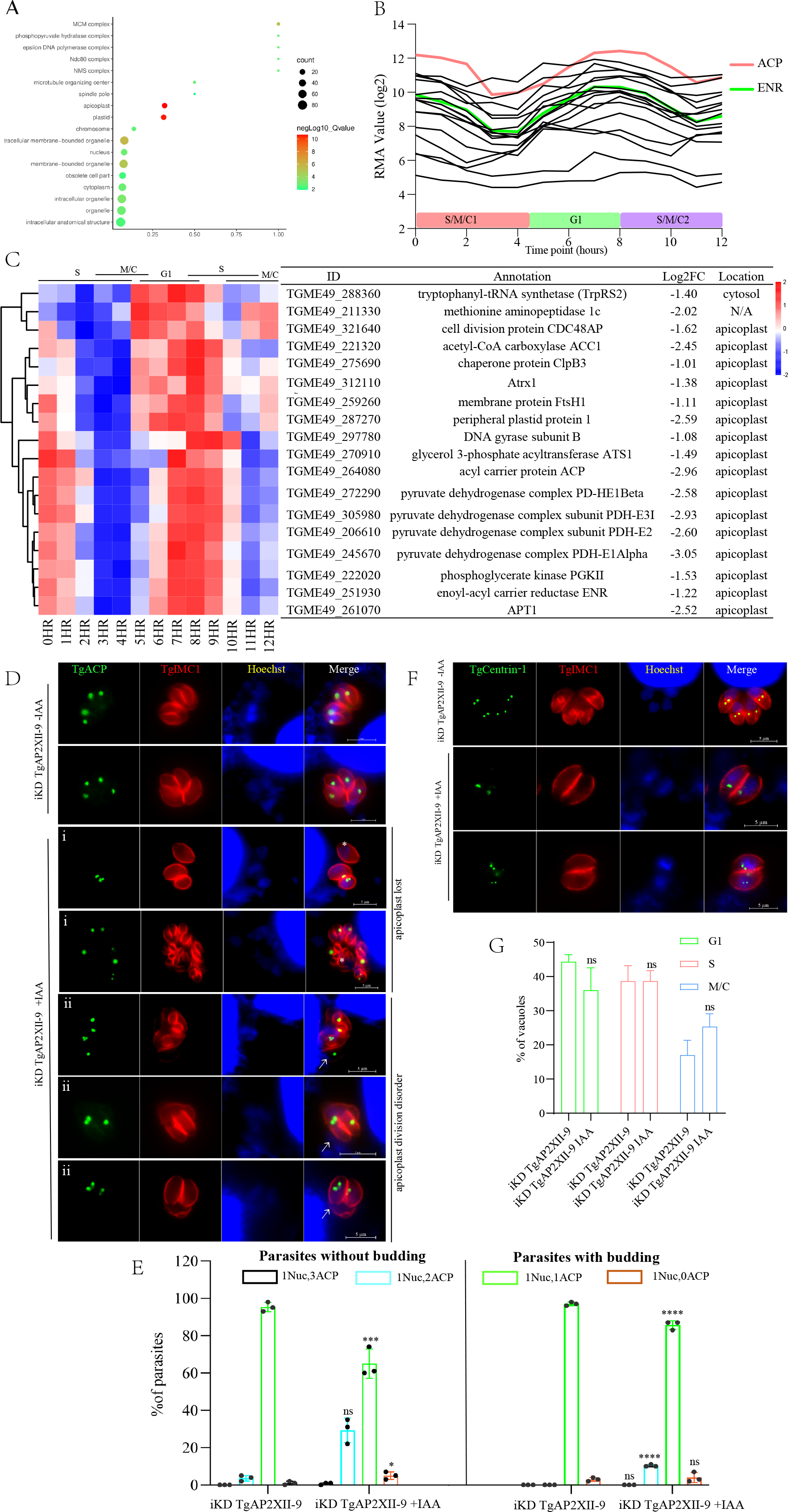
TgAP2XII-9 deficiency influences the expression of apicoplast genes. (A) GO analysis of cluster 1 protein in TgAP2XII-9-deficient parasites using ToxoDB annotations; significant GO terms are shown (Benjamini < 0.1). (B) Cell cycle expression patterns of 18 apicoplast genes down-regulated in TgAP2XII-9-deficient tachyzoites (9). RMA = robust multi-array average. (C) Heatmap showing down-regulation of 18 apicoplast genes after TgAP2XII-9 depletion. The putative annotation and location of these genes are listed. (D) Immunofluorescence analysis of apicoplast division in iKD TgAP2XII-9 parasites treated with or without IAA. Examples of different types of apicoplast abnormal divisions (shown by ACP staining) were observed. I) loss of apicoplast- asterisks mark parasites that produce 0 apicoplast; ii) apicoplast division disorder - arrows indicate 3 or 2 apicoplasts in a daughter cell and 1 apicoplast in another daughter cell. TgIMC1 (red) serves as a daughter cell budding marker. The nucleus was stained with Hoechst (bar = 5 µm). (E) Distribution of parasites containing the indicated number of segregated apicoplasts, as shown by Hoechst and TgACP staining on iKD TgAP2XII-9 parasites treated with or without IAA for 24 h. 1 Nuc indicates one nucleus; 0, 1, 2, and 3 ACP indicates loss of apicoplast, 1 apicoplast, 2 apicoplast, and 3 apicoplast, respectively. Parasites with or without daughter cell budding (determined by IMC1 staining) were plotted separately. 100 parasites with or without daughter cell budding were analyzed in each biological replicate. (F) Observation of centrosome division by IFA in iKD TgAP2XII-9 parasites treated with or without IAA. IFA was performed using antibodies to TgIMC1 (red) and TgCentrin1 (green). Hoechst was used to stain the nucleus. Parasites in the G1 phase contain single centrosomes, whereas parasites in the S-phase have duplicate centrosomes. The scale bar is shown on the lower right side of each image. (G) Quantification of phenotypes in 24 h-treated iKD TgAP2XII-9. At least 100 vacuoles were counted per biological replicate. Each experiment was repeated a total of 3 times. Results were shown as means ±SD of three independent experiments, and unpaired two-tailed Student’s *t*-test were employed. *** *P* < 0.001, **** *P* < 0.0001.

Two different types of unnatural divisions were mainly observed as follows: i) Loss of apicoplasts. Tachyzoites in the same PV exhibited both persistence and absence of apicoplast; ii) apicoplast division disorder. Tachyzoites in the same PV showed different numbers of apicoplast, including 0, 1, 2 and 3. Usually, the single copy apicoplast undergoes elongation and scission and divides equally into two daughter cells, but we observed disordered inheritance of apicoplast in the daughter cells. For instance, one daughter cell had two or three apicoplasts while the other had none (Fig. 6D). Quantification analysis demonstrated that 5% of vacuoles exhibited loss of apicoplasts after deletion of TgAP2XII-9 (Fig. 6E), and the frequency of vacuoles displaying two apicoplasts was significantly higher in a parasite with/without budding (Fig. 6E), with loss of apicoplast (1Nucl, 0 APC) also being noted. The same results were observed using the ENR antibody (Fig. S1A). The proportion of elongated apicoplast was also quantified; however, no significant difference was observed (Fig. S1B). Since segregation of apicoplasts into daughter parasites occurs only after centriole duplication (8), the division of the centriole was also observed by IFA using an anti-Centrin1 antibody. Besides, the results showed that the division of centriole was not affected after TgAP2XII-9 deletion (Fig. 6F and G). Additionally, the TgAP2XII-9 did not directly bind to the promoters of the apicoplast genes, as indicated from our CUT&Tag results. Thus, these results indicate that TgAP2XII-9 plays an indirect but crucial role in the accurate division and inheritance of apicoplast to daughter parasites.

Notably, a set of genes related to DNA duplication were downregulated upon TgAP2XII-9 deletion, with an expression pattern consistent with the cluster1 genes that are peaked in expression in the G1 phase (6-9 HR) (Fig. S2). These genes were also not affected by CUT&Tag enrichment, suggesting that TgAP2XII-9 may have an indirect role in regulating parasite DNA replication, but further validation is needed.

## DISCUSSION

*T. gondii* is an obligate intracellular apicomplexan parasite that infects almost all warm-blooded animals, including humans and livestock, and causes severe secondary infections in immunocompromised populations and reproductive disorders in pregnant women. It is estimated that about one-third of the global population is infected with *T. gondii* (43–45). *T. gondii* has a complex life cycle, including sexual and asexual stages. The tachyzoites can replicate rapidly by endodyogeny, and their cell cycle is divided into three phases (G1, S, and M phases) (6, 46). The rapid and smooth transformation of the parasite cell cycle relies on precise and complicated gene regulation; however, the mechanism of parasite cell cycle regulation is not fully understood. Twenty-four ApiAP2 transcription factors in *T. gondii* are predicted to be potentially involved in cell cycle regulation, including TgAP2XII-9. Here, we demonstrate that TgAP2XII-9 has a dynamic expression pattern during the parasite cell cycle, with peak expression in the S/M phase and a minimum level in the G1 phase. This study also shows that TgAP2XII-9 is essential for the growth of tachyzoites and that TgAP2XII-9 depletion leads to a significant repression of parasite invasion, replication and egress.

TgAP2XII-9 depletion also led to the downregulation of large amounts of cyclically expressed genes, including IMCs and ROPs/RONs, which are highly expressed in the S/M phase (Fig. 4E). Similarly, TgAP2X-5 deficiency resulted in the downregulation of genes highly expressed in the S/M phase (including multiple rhoptries and micronemes proteins), leading to a decrease in the virulence and invasive capacity of the parasite (7). Another study showed that knockdown of AP2X-4 resulted in the downregulation of several rhoptry proteins, especially those genes related to tissue cyst formation (ROP5, ROP17, ROP7, ROP2, ROP8 and ROP16), suggesting that AP2X-4 may affect parasitism by regulating these rhoptry proteins (16, 47). Interestingly, our results indicate that TgAP2XII-9 directly controls the expression of RON and ROP proteins (peak expression in S- and M- phase), including RON2, RON4, RON5 and RON 8, which are important components of the moving junction (26). MJ is a key structure for host cell invasion and is formed by intimate contact between the apical tip of the tachyzoite and the host cell membrane (26). The MJ of *T. gondii* tachyzoites consists of RON2, RON4, RON5, and RON8 secreted from rhoptry necks, and apical membrane antigen 1 (AMA1) secreted from micronemes (26). RON2 and RON5 are crucial for the invasion process of tachyzoites, and parasites lacking RON2 or RON5 cannot survive (48, 49). The invasive ability of RON8 and RON4-depleted parasites is significantly reduced by 70% and 60%, respectively (50, 51). In addition, AMA2 and AMA4 were found to be significantly downregulated in TgAP2XII-9-depleted parasites. AMA2 and AMA4 were found to be homologous to AMA1, but consist of a bradyzoite-specific MJ machinery and contribute to cyst burden (28). These results suggest that TgAP2XII-9 may affect the invasion of both tachyzoites and bradyzoites by regulating the transcription of MJ-related RON or AMA genes.

Notably, the transcription levels of a large number of IMC genes with peak expression in the S/M phases and minimum expression in the G1 phase are affected by TgAP2XII-9. IMC is a unique organelle that plays many important roles in the complex life cycle of apicomplexan parasites. IMC serves as a scaffold for daughter cell assembly and is crucial for maintaining the structural stability of tachyzoites (2, 52). Depletion of TgAP2XII-9 leads to the downregulation of 52 IMC proteins, some of which are known to be localized to daughter tachyzoites (35–39, 53). Interestingly, IMC29, IMC30, IMC31, IMC44, BCC3 and IMC36 were found to be localized only in the daughter buds of parasites, whereas AC12 and AC13 are located only in the apical cap of daughter buds (35). Of these, IMC29 has been proven to be an important early daughter bud component for replication, and parasites lacking IMC29 exhibit severe growth replication defects and loss of virulence (35). IMC subcompartmental proteins (ISPs) have been used to delineate the various sub compartments of the IMC through parasite division (37, 54). TgISP1 is present in the apical cap of both mother and daughter parasites, which is one of the first markers seen at initiation of daughter cell construction. TgISP2 and TgISP4 are localized in the central section of the IMC, and TgISP2 is considered to play an important role in regulating cell division in *T. gondii* (37, 52, 54, 55). In addition, after depletion of TgAP2XII-9, the transcription levels of cytoskeletal proteins expressed at the later stages of the replication process were also downregulated, including IMC1, IMC3, IMC4, IMC6, and IMC10. However, IMCs (IMC7, IMC12, IMC14, IMC37 and IMC40) expressed only in the mother cells were not affected (35, 56, 57). Subsequently, the disorganization exhibited by TgAP2XII-9-depleted parasites in daughter cell division was validated using IMC1 as the marker. However, not all of the above IMCs were directly controlled by TgAP2XII-9, but many of the directly controlled IMCs (IMC3, IMC4, IMC10, IMC31, IMC34, AC12, and AC13) were restricted by daughter cell formation or endodyogeny. In summary, these results suggest that TgAP2XII-9 may affect daughter division in tachyzoites by regulating the transcription of IMC genes related to daughter buds.

In addition, one of the important roles of the IMC is to act as an anchor for the actin-myosin motor complex which is necessary for both parasite invasion and egress (58, 59). GAP40 and GAP50 are glideosome-associated proteins that stably anchor the motor complex to the IMC (52, 60, 61), which is significantly downregulated in TgAP2XII-9 deficient parasites and is directly controlled by TgAP2XII-9. GAP50 is firmly immobilized in the IMC and is proposed to act as a fixed anchor for the motor complex (60, 62). GAP40 has been shown to interact with components of the motor complex and is expressed concurrently with GAP50 in early daughter cells, suggesting that GAP40 may also play a role in anchoring the motor complex (52, 60, 63). Therefore, in addition to ROP and RON proteins, the downregulation of the glideosome-associated proteins GAP40 and GAP50 may also be one of the reasons for the deficiency of parasite invasion and egress by TgAP2XII-9 knockdown.

TgAP2IX-5 has been shown to bind to the promoter of five ApiAP2s including TgAP2XII-9 and control a large number of IMC and apical complex genes (17). We compared the RNA-seq data of TgAP2IX-5 and TgAP2XII-9 and found that part of the IMCs were downregulated in TgAP2IX-5 or TgAP2XII-9 knockout parasites, including IMC1, IMC3, IMC6, IMC10, IMC29, IMC30, GAP40, AC2, AC4, AC7 and AC13 (17). Among these IMCs, the promoters of IMC3 and AC2 were shown to bind directly to TgAP2IX-5 and TgAP2XII-9 (17), which may indicate that TgAP2XII-9 not only acts as a downstream factor of TgAP2IX-5, but also co-opts with TgAP2IX-5 to activate the expression of IMC genes.

Recent studies have demonstrated that histone deacetylase (HDAC3) and microrchidia (MORC) interact with at least 12 ApiAP2s and act as a transcription repressor complex to suppress large amounts of genes (64). TgAP2XII-1 and TgAP2XI-2 have been shown to bind to the promoters of merozoite-specific genes as heterodimers and recruit MORC and HDAC3 to suppress these genes at the tachyzoite stage (64–67). The stage transition from tachyzoites to merozoites and the division transition from endodyogeny to endopolygeny were achieved by knockdown (KD) of these factors (64–67). Notably, double knockdown of TgAP2XII-1 and TgAP2XI-2 led to a near-complete stage transition (66, 67). In contrast, TgAP2XII-2 (another MORC-interacting ApiAP2) shows highly coordinated gene targets with MORC and HDAC3 (68, 69). However, its depletion activates only merogony gene expression, but not merogony or endopolygeny (67, 68). Apart from the MORC-HDAC3-ApiAP2s gene suppression complex, knockdown of cyclically regulated TgAP2IX-5 results in flexibility of cell cycle patterns from endodyogeny to endopolygeny (17). Chromatin immunoprecipitation results do not support direct binding of TgAP2IX-5 to merozoite-specific gene promoters but do support a large number of IMC genes and five ApiAP2s (TgAP2IV-4, TgAP2III-2, TgAP2XII-9, TgAP2X-9, and TgAP2XII-2) (19). Therefore, assuming a cascade ApiAP2 transcription regulation model and taking into account the unsuccessful inducement of merogony due to TgAP2XII-2 depletion (67), another key factor may exist in the remaining TgAP2IX-5 targeted to ApiAP2s.

In this study, small proportion of endopolygeny was also observed after TgAP2XII-9 knockout using TgIMC1 and TgGAP45 as staining markers. However, the abnormal divisions of daughter cells, including non-synchronous divisions and the presence of an odd number of tachyzoites in a PV, dominated the IAA-treated population. After TgAP2XII-9 knockout, we also found that 11% of DEGs were differently transcribed for enteroepithelial stage (EES, 996 in total) genes (70), but these genes showed both downregulation (97/208) and upregulation (111/208). In contrast, TgMORC depletion resulted in >80% EES gene upregulation (64), and TgAP2XII-1 and TgAP2XI-2 knockout also upregulated about half of the merozoite specific genes (65–67). Additionally, we stained TgAP2XII-9 knockout parasites with the merogony marker TgGRA11b, but observed negative signals (data not shown). Therefore, we speculate that TgAP2IX-5 controls the parasite division pattern from endodyogeny to endopolygeny through TgAP2XII-9, but not the stage transition from tachyzoites to merozoites as in the case of the MORC-HDAC3-AP2s complex dose (64–67).

Previous studies have revealed that subcellular organelles are strictly coordinated during tachyzoite replication, which invariably proceeds in the following order: centriole and Golgi → apicoplast → nucleus and ER → IMC; rhoptries and micronemes are synthesized de novo in each daughter cell (8). Our results indicate that deletion of TgAP2XII-9 results in significant downregulation of the transcriptional levels of many apicoplast genes, which have peak expression in G1 phase and lowest expression in the S/M phases. The peak expression of these apicoplast genes occurred significantly earlier than the IMC and ROP genes regulated by TgAP2XII-9, which is consistent with the previously reported theory that the division of apicoplasts occurs earlier than that of IMC and ROP during the division cycle of tachyzoites (8). This study subsequently analyzed the functions of these TgAP2XII-9-regulated apicoplast genes and found that half of these genes are key genes involved in the process of the apicoplast FASII pathway, including ACC1, ENR, APT1, ACP, ATS1, and the four PHDs (40, 41). FASII is a de novo fatty acid synthesis pathway restricted in the apicoplast of *T. gondii* and is essential throughout the tachyzoite life stage, as it provides the bulk of the fatty acids required for the synthesis of major membrane lipid classes (19, 40, 42). Dihydroxyacetone phosphate and phosphoenolpyruvate (PEP) were imported into the apicoplast through APT1 to provide fuel for the FASII pathway (19, 40, 71). The imported PEP was converted by the Pyruvate dehydrogenase complex into pyruvate and ultimately into acetyl CoA. Acetyl-CoA was then carboxylated by ACC1 to form malonyl-CoA (40, 41, 72, 73). Following the ACC1 activity, acetyl-CoA and malonyl-CoA were transferred to acyl carrier proteins, which transferred the nascent fatty acid chains to different enzymes in the FASII pathway (19, 40, 41, 72). A previous study showed that specific inhibition of the FASII pathway with triclosan affects apicoplast inheritance and parasite division by preventing cytokinesis completion, resulting in incomplete daughter cell budding (74). Multiple key components in the FASII pathway have been shown to be crucial for the formation of apicoplasts and the division of daughter parasites. ACP deficiency has been proven to result in defective apicoplast biogenesis and a consequent loss of the organelle (42). In addition, disruption of TgATS1 causes defects in organelle and daughter parasite development (40). We further confirmed that the apicoplast of the IFA-based TgAP2XII-9 parasites exhibits division disorder and organelle loss. Furthermore, abnormal budding and division of the daughter parasites were seen in TgAP2XII-9-deficient parasites (Fig. 5). These results indicate that TgAP2XII-9 deficiency leads to the downregulation of multiple apicoplast genes in the FAS II pathway, which in turn affects the division/formation of apicoplasts and further causes abnormal budding and division of tachyzoites.

## CONCLUSION

Our findings demonstrate that TgAP2XII-9 plays a pivotal role in activating the S/M-specific cell cycle program and influencing parasite invasion and division by directly targeting the MJ and IMC genes, and that TgAP2XII-9 knockout also disturbs apicoplast inheritance in *T. gondii* tachyzoites. This insight into asexual cell cycle regulation could help provide potential therapeutic targets and enhance our understanding of *T. gondii* cell cycle dynamics.

## MATERIALS AND METHODS

### Parasites and cell culture

*T. gondii* RH Δku80 strain and derivative strains were continuously cultured in vitro with human foreskin fibroblasts (HFFs; ATCC, Manassas, VA, USA) or African green monkey kidney cells (Vero cells, a gift from Prof Qun Liu, China Agricultural University) using Dulbecco’s Modified Eagle’s Medium (DMEM, Macgene, Beijing, China) with 2% c fetal bovine serum (FBS) (TransGen Biotech, Beijing, China) at 37°C and 5% CO2. Cells were cultured in DMEM supplemented with 10% FBS and incubated at 37℃ in a 5% CO2 environment.

### Generation of transgenic *T. gondii* strains

The EuPaGDT library was employed to design the corresponding guide RNAs for the gene-specific CRISPR-Cas9 plasmid used in this experiment. The construction of the CRISPR/Cas9 plasmid was performed as previously described(75). Briefly, Cas9 upstream and downstream fragments containing gRNA sequences were amplified and ligated with a seamless cloning kit (Vazyme Biotech, Co., Ltd, Nanjing). To construct iKD TgAP2XII-9 parasites, the mAID sequence and a 3× HA epitope tag were fused to the C-terminus of AP2XII-9. A plasmid (mAID-3HA-CAT-TIR1-3Flag) containing the chloramphenicol resistance gene (CmR) and the TIR1-3Flag expression cassette was constructed. For the C-terminal epitope tagging TgAP2XII-9, 59 bp PCR primers containing 39 bp fragments upstream of the TgAP2XII-9 translation stop codon and downstream of the gRNA site were designed to amplify PCR products from the mAID-3HA-CAT-TIR1-3Flag plasmid. The homologous recombination plasmid and the corresponding CRISPR–Cas9 plasmid were co-transfected into RH ΔKu80 parasites and screened in chloramphenicol-containing medium. Monoclonal parasites were identified using PCR and IFA. The degradation of AP2XII-9 was induced by IAA at a final concentration of 500 μM.

### Intracellular replication assay and invasion assay

The intracellular replication effect was observed by immunofluorescence microscopy. HFF cells in 12-well plates were infected with 1 × 10^5^ fresh RH tachyzoites per well and incubated for 1 h. The cell surface was then washed twice with PBS to remove extracellular tachyzoites, followed by incubation with IAA (500 μM) or vector (1:1000 ethanol). After 24 h of treatment, cells were fixed with 4% paraformaldehyde and then immunofluorescence staining was performed using rabbit anti-TgGAP45 antibodies (1:300, a gift from Professor Qun Liu in China Agricultural University) and Hoechst 33258 (1:100, Macgene, Beijing, China). The number of parasites per strain was determined by counting at least 100 vacuoles using fluorescence microscopy. For the invasion assay, the percentage of invasion was shown as the number of vacuoles per host cell. Three independent experiments were carried out.

### Egress Assay

Parasites were inoculated onto 12-well plates and cultured for 30 h with or without IAA treatment. The egress was triggered with 2 μM of Ca^2+^ ionophore A23187 (Macklin, Shanghai) for 2 min at 37℃ before fixation with PFA. The IFA was performed using rabbit anti-TgGAP45 antibodies. A total of 100 vacuoles were randomly selected to count ruptured vacuoles/whole vacuoles per slide. Three independent experiments were performed.

### Plaque assay

HFFs growing in 12-well plates were infected with 200 freshly harvested tachyzoites and incubated for 7 days without disturbance. The experimental group was incubated with vector (EtOH 1:1000) or IAA (500 μM). Thereafter, infected HFFs were fixed with 4% paraformaldehyde and visualized by staining with 0.2% crystal violet solution. The plaque area was counted by pixel using Photoshop C6S software (Adobe, USA), and data were compiled from three independent experiments.

### Immunofluorescence assay (IFA)

Freshly harvested parasites were inoculated onto HFF cells grown on glass coverslips in 12-well plates. After incubation, infected cells were properly fixed with 4% PFA for 1 h, permeabilized with 0.25% Triton X-100, and then blocked with 3% bovine serum albumin (BSA) for 30 min. The samples were incubated with mouse anti-HA (1:500, Sigma, USA), rabbit anti-TgGAP45 (1:300, a gift from Professor Qun Liu in China Agricultural University), rabbit anti-TgIMC1 antibody (1:300, an inner membrane complex marker), mouse anti-TgACP (1:300, a marker for apicoplast, from Prof. Liu at China Agricultural University), mouse anti-ENR (1:100, from Prof. Liu at China Agricultural University) and rabbit anti-αCentrin 1 antibody (1:500, a gift from Professor Shaojun Long in China Agricultural University) for 1 h and washed three times with PBS, followed by incubation with secondary FITC- or cy3-conjugated antibodies (1:100, Proteintech, USA) for 1 h. Images were obtained using a Zeiss Fluorescence Microscopy system (Zeiss, Germany).

### TgAP2XII-9 recovery experiments

Parasites of iKD TgAP2XII-9 strain were inoculated on HFF cells grown on coverslips of a 12-well plate in the presence of auxin for 16 h. Auxin was washed off and parasites were allowed to grow in IAA-free medium for additional 0, 3, 6, and 20 h before being fixed with 4% PFA. Parasite nuclei were labelled with hoechst33258 and the parasite’s inner membrane complex (IMC) was labelled with TgIMC1. Three independent experiments were performed.

### RNA-Seq and data analysis

Transgenic parasite strains cultured in Vero cells were treated with either 500 μM IAA or vehicle for 24 h. Total RNAs from *T. gondii* tachyzoites were then extracted using the M5 Total RNA Extraction Reagent (Mei5 Biotechnology Co., Ltd, Beijing) according to the manufacturer’s protocol. Each treatment consisted of three biological replicates. The purity, concentration and integrity of RNAs were tested using the NanoPhotometer® (IMPLEN, CA, USA), the Qubit® RNA Assay Kit in Qubit® 2.0 Fluorometer (Life Technologies, CA, USA) and the RNA Nano 6000 Assay Kit of the Bioanalyzer 2100 system (Agilent Technologies, CA, USA), respectively. Only qualified samples were used for library preparation. Illumina sequencing libraries were generated using the NEBNext® Ultra™ RNA Library Prep Kit for Illumina® (NEB, USA) according to the manufacturer’s recommendations. Sequencing was performed using the Illumina Novaseq 6000 platform from Shanghai Personal Biotechnology Co., Ltd. to generate 150 bp paired-end reads. The original sequencing data can be found in the Sequence Read Archive database under the accession number PRJNA1070975.

RNA-seq clean reads were uploaded to the BMKCloud (www.biocloud.net) platform for analysis based on the reference genome of Toxoplasma Type II ME49 strain (ToxoDB-57). Briefly, Paired-end clean reads were aligned to the reference genome using Hisat2 (76). Read counts were calculated for each gene using the sorted bam files from StringTie (77). Differentially expressed genes (DEGs) between treated and untreated parasites were calculated by edgeR (78). Gene ontology enrichment analysis was performed in ToxoDB. Gene expression with a fold change >2 or < -2 and FDR < 0.05 was considered as significantly differentially expressed. TPM (Transcripts per kilobase million) values were calculated for each gene and used for generating clustered heatmaps. Gene clustering was performed using Mfuzz from the R package. Gene Ontology enrichment analysis was conducted utilizing topGO (79) and GO annotations accessible on ToxoDB.org (version 64), with significant GO terms identified based on a Benjamini < 0.1.

### CUT&Tag and data analysis

Transgenic strains were inoculated in 25 cm^2^ cell culture flasks and washed after 4 h to remove non-invasive parasites. Transgenic strains (1 x 10^7^) were harvested after 20 h of incubation. Library construction was performed using the NovoNGS CUT&Tag 4.0 High-Sensitivity Kit for Illumina (Novoprotein, Suzhou, China) according to the manufacturer’s instructions. Briefly, fresh tachyzoites were bound to activated concanavalin A beads (10 μL/sample) and incubated for 10 min at room temperature. The mixture was resuspended and incubated with primary antibody (1:50, mouse anti-HA) at 4°C overnight. After several washes, the parasites were incubated with secondary antibody (1:100, goat anti-mouse IgG) for 1 h at room temperature. The parasites were then resuspended with pAG-Transposome buffer and incubated for 1 h at room temperature on a rotator. Tagmentation was stopped by MgCl2 treatment and DNA extraction was performed using DNA extract beads (Novoprotein, Suzhou, China). Illumina sequencing libraries were generated by PCR amplification using specific adaptors according to the manufacturer’s recommendations (NovoNGS CUT&Tag 4.0 High-Sensitivity Kit for Illumina B box, Novoprotein, Suzhou, China). CUT&Tag libraries were sequenced using the Illumina Novaseq 6000 platform (Beijing Novogene Technology Co., Ltd).

The paired-end reads were filtered and then aligned to the *T. gondii* ME49 reference genome using Bowtie2 (80) (v.2.1.0). The resulting sam files were transformed into bam files. PCR duplicates were removed from the sorted bam files using Picard tools (https://broadinstitute.github.io/picard/). The filtered reads were then employed to identify CUT&Tag peaks using MACS2 (81). The overlapped peaks in the two biological replicates were identified by the Irreproducibility Discovery Rate (IDR) (82). Final peaks were annotated against the latest *T. gondii* reference genome in ToxoDB. The sorted and filtered bam files of CUT&Tag peaks and RNA-seq reads were normalized to RPKM with a resolution of 10 bp (bin size) and transformed into bigwig files for direct visualization in IGV (Integrative Genomics Viewer) (83). Raw sequencing data and processed data are available in the NCBI GEO database under the accession number GSE266204.

### Statistical analysis

Violin charts, line drawings, scatter plots, and histograms were generated using GraphPad Prism 9 (San Diego, CA, USA). Heatmaps were drawn using the OmicStudio tool on https://www.omicstudio.cn/tool. Time series analysis was performed online in BioLadder (bioladder.cn). All experiments were performed in independent biological replicates as described for each experiment in the manuscript. Statistical significance in plaque assay, invasion, proliferation, and parasite growth inhibition assay was evaluated by two-tailed unpaired *t*-tests or two-way ANOVA using GraphPad Prism. Statistical data are expressed as mean value ± standard error.

## ACKNOWLEDGMENTS

We are grateful to Prof. Qun Liu, Prof. Jing Liu and Prof. Shaojun Long (China Agricultural University, China) for providing parasite strains and antibodies. Prof. Dongying Wang, Mao Huang, Xinru Cao and Yazhen Ma in our institution are also acknowledged for their technical help.

## AUTHOR CONTRIBUTIONS

Xingju Song. and Dandan Hu conceived and designed the study. Yuehong Shi performed the experiments and analyzed the data. Xuan Li and Yingying Xue helped in validation and visualization. Xingju Song and Yuehong Shi drafted the manuscript. Dandan Hu and Xingju Song funded, reviewed and edited the manuscript. All authors read and approved the final manuscript.

## DATA AVAILABILITY

RNA-Seq data generated in this study have been deposited to the NCBI Sequence Archive (SRA) under the accession numbers PRJNA1070975. Raw sequencing data and processed data for CUT&Tag experiments are available in the NCBI GEO database under the accession number GSE266204.

## SUPPLEMENTARY MATERIALS

Figure S1. **TgAP2XII-9 deficiency affects the division of apicoplast.** (A) IFA shows the apicoplast division (shown by ENR staining) in iKD TgAP2XII-9 parasites treated with or without IAA. Patterns of ENR were expressed with/without TgAP2XII-9. i) loss of apicoplast; ii) apicoplast division disorder. Asterisks mark parasites producing 0 apicoplast; arrowheads are pointed to a single maternal parasite or a daughter bud producing 2 apicoplast, which indicates dysregulation of apicoplast replication. TgIMC1 (green) serves as a daughter cell budding marker. The nucleus was stained with Hoechst (bar = 5 µm). (B) Bar graph representing the percentage of iKD TgAP2XII-9 parasites with an elongated plastid in the absence and presence of auxin treatment for 24 h. A two-tailed Student’s *t*-test was performed: \**P* < 0.05. (n=3 independent experiments)

Figure S2. **TgAP2XII-9 regulates DNA replication in tachyzoites.** (A) KEGG pathway enrichment of differentially expressed genes (DEGs). The X-axis shows the rich factor and the Y-axis corresponds to the KEGG pathway. The size of the dots represents the number of DEGs. The colors of the dots represent the enrichment-adjusted P-values. (B) Expression of transcripts related to DNA replication after deletion of TgAP2XII-9 (9) (data source: ToxoDB.org); RMA, robust multiarray average; G1, G1 phase; S, synthesis phase; M, mitosis phase; C, cytokinesis phase. (C) Heatmap displaying the cell cycle expression profile of 16 genes related to the down-regulation of DNA replication in TgAP2XII-9 defective tachyzoites (9). The mean log 2 FPKM value for every gene in each group was normalized and used. The approximate timing of the budding cycle is indicated at the top.

Table S1. Primers used in this study.

Table S2. Integrated CUT&Tag hits and transcariptomic data for the iKD TgAP2XII-9 strain.

## REFERENCES

1. Montoya JG and Liesenfeld O. 2004. Toxoplasmosis. Lancet 363(9425):1965–1976.

2. Blader IJ, Coleman BI, Chen CT and Gubbels MJ. 2015. Lytic Cycle of Toxoplasma gondii: 15 Years Later. Annu Rev Microbiol 69:463–485.

3. Dubey JP. 2009. History of the discovery of the life cycle of Toxoplasma gondii. Int J Parasitol 39(8):877–882.

4. Francia ME and Striepen B. 2014. Cell division in apicomplexan parasites. Nat Rev Microbiol 12(2):125–136.

5. Gubbels MJ, White M and Szatanek T. 2008. The cell cycle and Toxoplasma gondii cell division: tightly knit or loosely stitched? Int J Parasitol 38(12):1343–1358.

6. Kim K. 2018. The Epigenome, Cell Cycle, and Development in Toxoplasma. Annu Rev Microbiol 72:479–499.

7. Lesage KM, Huot L, Mouveaux T, Courjol F, Saliou JM and Gissot M. 2018. Cooperative binding of ApiAP2 transcription factors is crucial for the expression of virulence genes in Toxoplasma gondii. Nucleic Acids Res 46(12):6057–6068.

8. Nishi M, Hu K, Murray JM and Roos DS. 2008. Organellar dynamics during the cell cycle of Toxoplasma gondii. J Cell Sci 121(Pt 9):1559–1568.

9. Behnke MS, Wootton JC, Lehmann MM, Radke JB, Lucas O, Nawas J, Sibley LD and White MW. 2010. Coordinated progression through two subtranscriptomes underlies the tachyzoite cycle of Toxoplasma gondii. PLoS One 5(8):e12354.

10. Balaji S, Babu MM, Iyer LM and Aravind L. 2005. Discovery of the principal specific transcription factors of Apicomplexa and their implication for the evolution of the AP2-integrase DNA binding domains. Nucleic Acids Res 33(13):3994–4006.

11. Walker R, Gissot M, Croken MM, Huot L, Hot D, Kim K and Tomavo S. 2013. The Toxoplasma nuclear factor TgAP2XI-4 controls bradyzoite gene expression and cyst formation. Mol Microbiol 87(3):641–655.

12. Huang S, Holmes MJ, Radke JB, Hong DP, Liu TK, White MW and Sullivan WJ, Jr. 2017. Toxoplasma gondii AP2IX-4 Regulates Gene Expression during Bradyzoite Development. mSphere 2(2).

13. Hong DP, Radke JB and White MW. 2017. Opposing Transcriptional Mechanisms Regulate Toxoplasma Development. mSphere 2(1).

14. Radke JB, Worth D, Hong D, Huang S, Sullivan WJ, Jr., Wilson EH and White MW. 2018. Transcriptional repression by ApiAP2 factors is central to chronic toxoplasmosis. PLoS Pathog 14(5):e1007035.

15. Radke JB, Lucas O, De Silva EK, Ma Y, Sullivan WJ, Jr., Weiss LM, Llinas M and White MW. 2013. ApiAP2 transcription factor restricts development of the Toxoplasma tissue cyst. Proc Natl Acad Sci U S A 110(17):6871–6876.

16. Zhang J, Fan F, Zhang L and Shen B. 2022. Nuclear Factor AP2X-4 Governs the Expression of Cell Cycle- and Life Stage-Regulated Genes and is Critical for Toxoplasma Growth. Microbiol Spectr 10(4):e0012022.

17. Khelifa AS, Guillen Sanchez C, Lesage KM, Huot L, Mouveaux T, Pericard P, Barois N, Touzet H, Marot G, Roger E and Gissot M. 2021. TgAP2IX-5 is a key transcriptional regulator of the asexual cell cycle division in Toxoplasma gondii. Nat Commun 12(1):116.

18. Wang C, Hu D, Tang X, Song X, Wang S, Zhang S, Duan C, Sun P, Suo J, Ma H, Suo X and Liu X. 2021. Internal daughter formation of Toxoplasma gondii tachyzoites is coordinated by transcription factor TgAP2IX-5. Cell Microbiol 23(3):e13291.

19. Ramakrishnan S, Docampo MD, Macrae JI, Pujol FM, Brooks CF, van Dooren GG, Hiltunen JK, Kastaniotis AJ, McConville MJ and Striepen B. 2012. Apicoplast and endoplasmic reticulum cooperate in fatty acid biosynthesis in apicomplexan parasite Toxoplasma gondii. J Biol Chem 287(7):4957–4971.

20. Radke JB, Worth D, Hong D, Huang S, Sullivan Jr WJ, Wilson EH and White MWJPp. 2018. Transcriptional repression by ApiAP2 factors is central to chronic toxoplasmosis. 14(5):e1007035.

21. Srivastava S, White MW and Sullivan Jr WJJM. 2020. Toxoplasma gondii AP2XII-2 contributes to proper progression through S-phase of the cell cycle. 5(5):10.1128/msphere.00542-00520.

22. Srivastava S, Holmes MJ, White MW and Sullivan Jr WJJM. 2023. Toxoplasma gondii AP2XII-2 contributes to transcriptional repression for sexual commitment. 8(2):e00606-00622.

23. Hawkins LM, Wang C, Chaput D, Batra M, Marsilia C, Awshah D and Suvorova ESJb. 2023. The G2 phase controls binary division of Toxoplasma gondii.2023.2007. 2031.551351.

24. Jeninga MD, Quinn JE and Petter M. 2019. ApiAP2 Transcription Factors in Apicomplexan Parasites. Pathogens 8(2).

25. Altschul SF, Wootton JC, Zaslavsky E and Yu YK. 2010. The construction and use of log-odds substitution scores for multiple sequence alignment. PLoS Comput Biol 6(7):e1000852.

26. Besteiro S, Dubremetz JF and Lebrun M. 2011. The moving junction of apicomplexan parasites: a key structure for invasion. Cell Microbiol 13(6):797–805.

27. Lamarque MH, Roques M, Kong-Hap M, Tonkin ML, Rugarabamu G, Marq J-B, Penarete-Vargas DM, Boulanger MJ, Soldati-Favre D and Lebrun MJNc. 2014. Plasticity and redundancy among AMA–RON pairs ensure host cell entry of Toxoplasma parasites. 5(1):4098.

28. Najm R, Ruivo MTG, Penarete-Vargas DM, Hamie M, Mouveaux T, Gissot M, Boulanger MJ, El Hajj H and Lebrun MJPotNAoS. 2023. Invasion of Toxoplasma gondii bradyzoites: Molecular dissection of the moving junction proteins and effective vaccination targets. 120(5):e2219533120.

29. Butterworth S, Torelli F, Lockyer EJ, Wagener J, Song O-R, Broncel M, Russell MR, Moreira-Souza ACA, Young JC and Treeck MJPP. 2022. Toxoplasma gondii virulence factor ROP1 reduces parasite susceptibility to murine and human innate immune restriction. 18(12):e1011021.

30. Khaminets A, Hunn JP, Könen-Waisman S, Zhao YO, Preukschat D, Coers J, Boyle JP, Ong YC, Boothroyd JC and Reichmann GJCm. 2010. Coordinated loading of IRG resistance GTPases on to the Toxoplasma gondii parasitophorous vacuole. 12(7):939–961.

31. Etheridge RD, Alaganan A, Tang K, Lou HJ, Turk BE, Sibley LDJCh and microbe. 2014. The Toxoplasma pseudokinase ROP5 forms complexes with ROP18 and ROP17 kinases that synergize to control acute virulence in mice. 15(5):537–550.

32. Singh S, Murillo-León M, Endres NS, Arenas Soto AF, Gómez-Marín JE, Melbert F, Kanneganti T-D, Yamamoto M, Campos C and Howard JCJPP. 2023. ROP39 is an Irgb10-specific parasite effector that modulates acute Toxoplasma gondii virulence. 19(1):e1011003.

33. Camejo A, Gold DA, Lu D, McFetridge K, Julien L, Yang N, Jensen KD and Saeij JPJIjfp. 2014. Identification of three novel Toxoplasma gondii rhoptry proteins. 44(2):147–160.

34. Tosetti N, Dos Santos Pacheco N, Bertiaux E, Maco B, Bournonville L, Hamel V, Guichard P and Soldati-Favre D. 2020. Essential function of the alveolin network in the subpellicular microtubules and conoid assembly in Toxoplasma gondii. Elife 9.

35. Back PS, Moon AS, Pasquarelli RR, Bell HN, Torres JA, Chen AL, Sha J, Vashisht AA, Wohlschlegel JA and Bradley PJ. 2023. IMC29 Plays an Important Role in Toxoplasma Endodyogeny and Reveals New Components of the Daughter-Enriched IMC Proteome. mBio 14(1):e0304222.

36. Pasquarelli RR, Back PS, Sha J, Wohlschlegel JA and Bradley PJ. 2023. Identification of IMC43, a novel IMC protein that collaborates with IMC32 to form an essential daughter bud assembly complex in Toxoplasma gondii. PLoS Pathog 19(10):e1011707.

37. Beck JR, Rodriguez-Fernandez IA, de Leon JC, Huynh MH, Carruthers VB, Morrissette NS and Bradley PJ. 2010. A novel family of Toxoplasma IMC proteins displays a hierarchical organization and functions in coordinating parasite division. PLoS Pathog 6(9):e1001094.

38. Vigetti L, Labouré T, Roumégous C, Cannella D, Touquet B, Mayer C, Couté Y, Frénal K, Tardieux I and Renesto P. 2022. The BCC7 Protein Contributes to the Toxoplasma Basal Pole by Interfacing between the MyoC Motor and the IMC Membrane Network. Int J Mol Sci 23(11).

39. Mann T, Gaskins E and Beckers C. 2002. Proteolytic processing of TgIMC1 during maturation of the membrane skeleton of Toxoplasma gondii. J Biol Chem 277(43):41240–41246.

40. Amiar S, MacRae JI, Callahan DL, Dubois D, van Dooren GG, Shears MJ, Cesbron-Delauw MF, Maréchal E, McConville MJ, McFadden GI, Yamaryo-Botté Y and Botté CY. 2016. Apicoplast-Localized Lysophosphatidic Acid Precursor Assembly Is Required for Bulk Phospholipid Synthesis in Toxoplasma gondii and Relies on an Algal/Plant-Like Glycerol 3-Phosphate Acyltransferase. PLoS Pathog 12(8):e1005765.

41. Coppens I and Botté C Biochemistry and metabolism of Toxoplasma gondii: lipid synthesis and uptake. in: Toxoplasma gondii (Ed.^, Eds.), Elsevier, 2020, 367-395.

42. Mazumdar J, E HW, Masek K, C AH and Striepen B. 2006. Apicoplast fatty acid synthesis is essential for organelle biogenesis and parasite survival in Toxoplasma gondii. Proc Natl Acad Sci U S A 103(35):13192–13197.

43. Montoya JG LO. 2004. Toxoplasmosis. Lancet 363(9425):1965–1976.

44. Nayeri T, Sarvi S, Moosazadeh M, Amouei A, Hosseininejad Z and Daryani A. 2020. The global seroprevalence of anti-Toxoplasma gondii antibodies in women who had spontaneous abortion: A systematic review and meta-analysis. PLoS neglected tropical diseases 14(3):e0008103.

45. Wang ZD, Wang SC, Liu HH, Ma HY, Li ZY, Wei F, Zhu XQ and Liu Q. 2017. Prevalence and burden of Toxoplasma gondii infection in HIV-infected people: a systematic review and meta-analysis. Lancet HIV 4(4):e177–e188.

46. Radke JR, Striepen B, Guerini MN, Jerome ME, Roos DS and White MW. 2001. Defining the cell cycle for the tachyzoite stage of Toxoplasma gondii. Mol Biochem Parasitol 115(2):165–175.

47. Fox BA, Rommereim LM, Guevara RB, Falla A, Hortua Triana MA, Sun Y and Bzik DJ. 2016. The Toxoplasma gondii Rhoptry Kinome Is Essential for Chronic Infection. mBio 7(3).

48. Beck JR, Chen AL, Kim EW and Bradley PJ. 2014. RON5 is critical for organization and function of the Toxoplasma moving junction complex. PLoS Pathog 10(3):e1004025.

49. Lamarque MH, Roques M, Kong-Hap M, Tonkin ML, Rugarabamu G, Marq JB, Penarete-Vargas DM, Boulanger MJ, Soldati-Favre D and Lebrun M. 2014. Plasticity and redundancy among AMA-RON pairs ensure host cell entry of Toxoplasma parasites. Nat Commun 5:4098.

50. Straub KW, Peng ED, Hajagos BE, Tyler JS and Bradley PJ. 2011. The moving junction protein RON8 facilitates firm attachment and host cell invasion in Toxoplasma gondii. PLoS Pathog 7(3):e1002007.

51. Guérin A, Corrales RM, Parker ML, Lamarque MH, Jacot D, El Hajj H, Soldati-Favre D, Boulanger MJ and Lebrun M. 2017. Efficient invasion by Toxoplasma depends on the subversion of host protein networks. Nat Microbiol 2(10):1358–1366.

52. Harding CR and Meissner M. 2014. The inner membrane complex through development of Toxoplasma gondii and Plasmodium. Cell Microbiol 16(5):632–641.

53. Engelberg K, Bechtel T, Michaud C, Weerapana E and Gubbels MJ. 2022. Proteomic characterization of the Toxoplasma gondii cytokinesis machinery portrays an expanded hierarchy of its assembly and function. Nat Commun 13(1):4644.

54. Poulin B, Patzewitz EM, Brady D, Silvie O, Wright MH, Ferguson DJ, Wall RJ, Whipple S, Guttery DS, Tate EW, Wickstead B, Holder AA and Tewari R. 2013. Unique apicomplexan IMC sub-compartment proteins are early markers for apical polarity in the malaria parasite. Biol Open 2(11):1160–1170.

55. Fung C, Beck JR, Robertson SD, Gubbels MJ and Bradley PJ. 2012. Toxoplasma ISP4 is a central IMC sub-compartment protein whose localization depends on palmitoylation but not myristoylation. Mol Biochem Parasitol 184(2):99–108.

56. Anderson-White BR, Ivey FD, Cheng K, Szatanek T, Lorestani A, Beckers CJ, Ferguson DJ, Sahoo N and Gubbels MJ. 2011. A family of intermediate filament-like proteins is sequentially assembled into the cytoskeleton of Toxoplasma gondii. Cell Microbiol 13(1):18–31.

57. Dubey R, Harrison B, Dangoudoubiyam S, Bandini G, Cheng K, Kosber A, Agop-Nersesian C, Howe DK, Samuelson J, Ferguson DJP and Gubbels MJ. 2017. Differential Roles for Inner Membrane Complex Proteins across Toxoplasma gondii and Sarcocystis neurona Development. mSphere 2(5).

58. Opitz C and Soldati D. 2002. ’The glideosome’: a dynamic complex powering gliding motion and host cell invasion by Toxoplasma gondii. Mol Microbiol 45(3):597–604.

59. Frénal K, Dubremetz JF, Lebrun M and Soldati-Favre D. 2017. Gliding motility powers invasion and egress in Apicomplexa. Nat Rev Microbiol 15(11):645–660.

60. Frénal K, Polonais V, Marq JB, Stratmann R, Limenitakis J and Soldati-Favre D. 2010. Functional dissection of the apicomplexan glideosome molecular architecture. Cell Host Microbe 8(4):343–357.

61. Morrissette N and Gubbels M-J. 2014. The Toxoplasma cytoskeleton: structures, proteins and processes. Toxoplasma gondii:455–503.

62. Johnson TM, Rajfur Z, Jacobson K and Beckers CJ. 2007. Immobilization of the type XIV myosin complex in Toxoplasma gondii. Mol Biol Cell 18(8):3039–3046.

63. Fauquenoy S, Hovasse A, Sloves PJ, Morelle W, Dilezitoko Alayi T, Slomianny C, Werkmeister E, Schaeffer C, Van Dorsselaer A and Tomavo S. 2011. Unusual N-glycan structures required for trafficking Toxoplasma gondii GAP50 to the inner membrane complex regulate host cell entry through parasite motility. Mol Cell Proteomics 10(9):M111.008953.

64. Farhat DC, Swale C, Dard C, Cannella D, Ortet P, Barakat M, Sindikubwabo F, Belmudes L, De Bock PJ, Couté Y, Bougdour A and Hakimi MA. 2020. A MORC-driven transcriptional switch controls Toxoplasma developmental trajectories and sexual commitment. Nat Microbiol 5(4):570–583.

65. Fan F, Xue L, Yin X, Gupta N and Shen B. 2023. AP2XII-1 is a negative regulator of merogony and presexual commitment in Toxoplasma gondii. mBio:e0178523.

66. Wang JL, Li TT, Zhang NZ, Wang M, Sun LX, Zhang ZW, Fu BQ, Elsheikha HM and Zhu XQ. 2024. The transcription factor AP2XI-2 is a key negative regulator of Toxoplasma gondii merogony. Nat Commun 15(1):793.

67. Antunes AV, Shahinas M, Swale C, Farhat DC, Ramakrishnan C, Bruley C, Cannella D, Robert MG, Corrao C, Couté Y, Hehl AB, Bougdour A, Coppens I and Hakimi MA. 2024. In vitro production of cat-restricted Toxoplasma pre-sexual stages. Nature 625(7994):366–376.

68. Srivastava S, Holmes MJ, White MW and Sullivan WJ, Jr. 2023. Toxoplasma gondii AP2XII-2 Contributes to Transcriptional Repression for Sexual Commitment. mSphere 8(2):e0060622.

69. Srivastava S, White MW and Sullivan WJ, Jr. 2020. Toxoplasma gondii AP2XII-2 Contributes to Proper Progression through S-Phase of the Cell Cycle. mSphere 5(5).

70. Ramakrishnan C, Maier S, Walker RA, Rehrauer H, Joekel DE, Winiger RR, Basso WU, Grigg ME, Hehl AB and Deplazes PJSr. 2019. An experimental genetically attenuated live vaccine to prevent transmission of Toxoplasma gondii by cats. 9(1):1474.

71. Lim L, Kalanon M and McFadden GI. 2009. New proteins in the apicoplast membranes: time to rethink apicoplast protein targeting. Trends Parasitol 25(5):197–200.

72. Dubois D, Fernandes S, Amiar S, Dass S, Katris NJ, Botté CY and Yamaryo-Botté Y. 2018. Toxoplasma gondii acetyl-CoA synthetase is involved in fatty acid elongation (of long fatty acid chains) during tachyzoite life stages. J Lipid Res 59(6):994–1004.

73. Jelenska J, Crawford MJ, Harb OS, Zuther E, Haselkorn R, Roos DS and Gornicki P. 2001. Subcellular localization of acetyl-CoA carboxylase in the apicomplexan parasite Toxoplasma gondii. Proc Natl Acad Sci U S A 98(5):2723–2728.

74. Martins-Duarte É S, Carias M, Vommaro R, Surolia N and de Souza W. 2016. Apicoplast fatty acid synthesis is essential for pellicle formation at the end of cytokinesis in Toxoplasma gondii. J Cell Sci 129(17):3320–3331.

75. Fu Y, Cui X, Fan S, Liu J, Zhang X, Wu Y and Liu Q. 2018. Comprehensive Characterization of Toxoplasma Acyl Coenzyme A-Binding Protein TgACBP2 and Its Critical Role in Parasite Cardiolipin Metabolism. mBio 9(5).

76. Kim D, Paggi JM, Park C, Bennett C and Salzberg SL. 2019. Graph-based genome alignment and genotyping with HISAT2 and HISAT-genotype. Nat Biotechnol 37(8):907–915.

77. Pertea M, Kim D, Pertea GM, Leek JT and Salzberg SL. 2016. Transcript-level expression analysis of RNA-seq experiments with HISAT, StringTie and Ballgown. Nat Protoc 11(9):1650–1667.

78. Robinson MD, McCarthy DJ and Smyth GK. 2010. edgeR: a Bioconductor package for differential expression analysis of digital gene expression data. Bioinformatics 26(1):139–140.

79. Alexa A, Rahnenführer J and Lengauer T. 2006. Improved scoring of functional groups from gene expression data by decorrelating GO graph structure. Bioinformatics 22(13):1600–1607.

80. Langmead B and Salzberg SL. 2012. Fast gapped-read alignment with Bowtie 2. Nat Methods 9(4):357–359.

81. Yashar WM, Kong G, VanCampen J, Curtiss BM, Coleman DJ, Carbone L, Yardimci GG, Maxson JE and Braun TP. 2022. GoPeaks: histone modification peak calling for CUT&Tag. Genome biology 23(1):144.

82. Singh R, Zhang F and Li Q. 2022. Assessing reproducibility of high-throughput experiments in the case of missing data. Stat Med 41(10):1884–1899.

83. Thorvaldsdóttir H, Robinson JT and Mesirov JP. 2013. Integrative Genomics Viewer (IGV): high-performance genomics data visualization and exploration. Brief Bioinform 14(2):178–192.

