## Supplementary figures and images for "Cyclical transcription factor AP2XII-9 is a key activator for asexual division and apicoplast inheritance in *Toxoplasma gondii* tachyzoite"

### Fig. S1

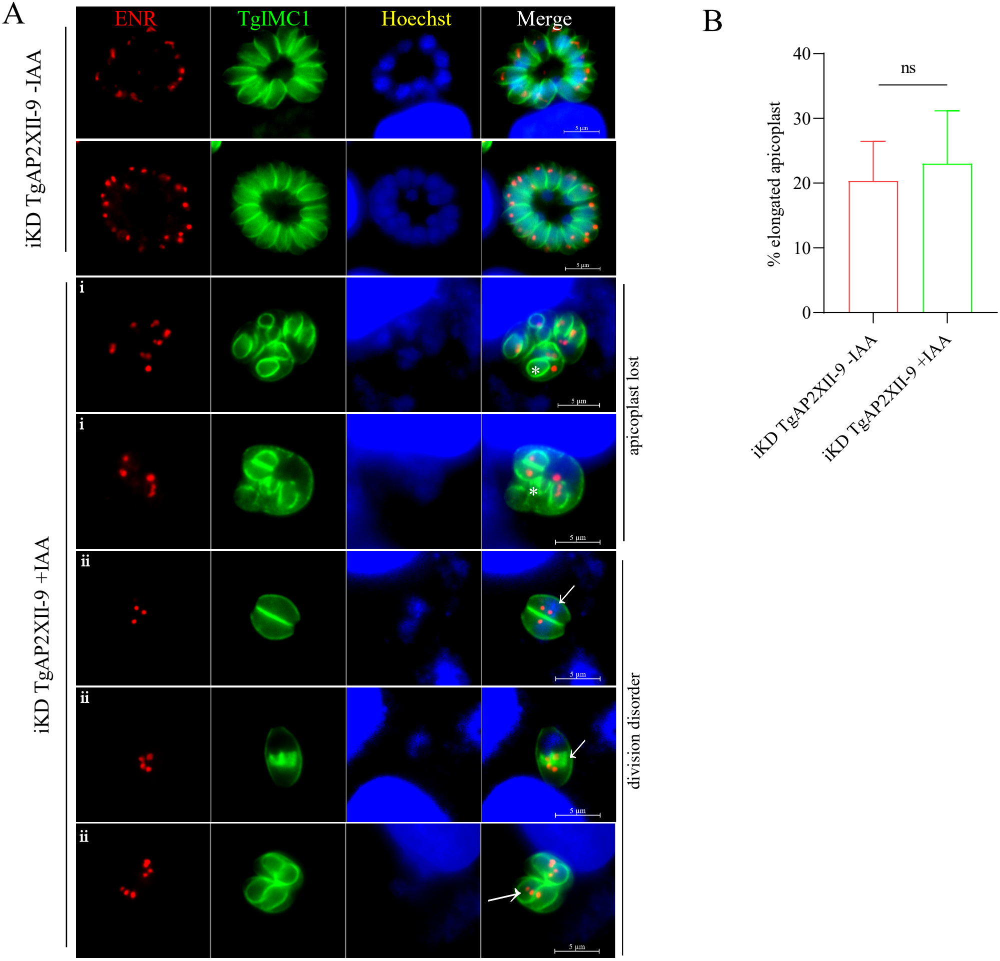
