## Supplementary material for "Cyclical transcription factor AP2XII-9 is a key activator for asexual division and apicoplast inheritance in *Toxoplasma gondii* tachyzoite": Fig. S2

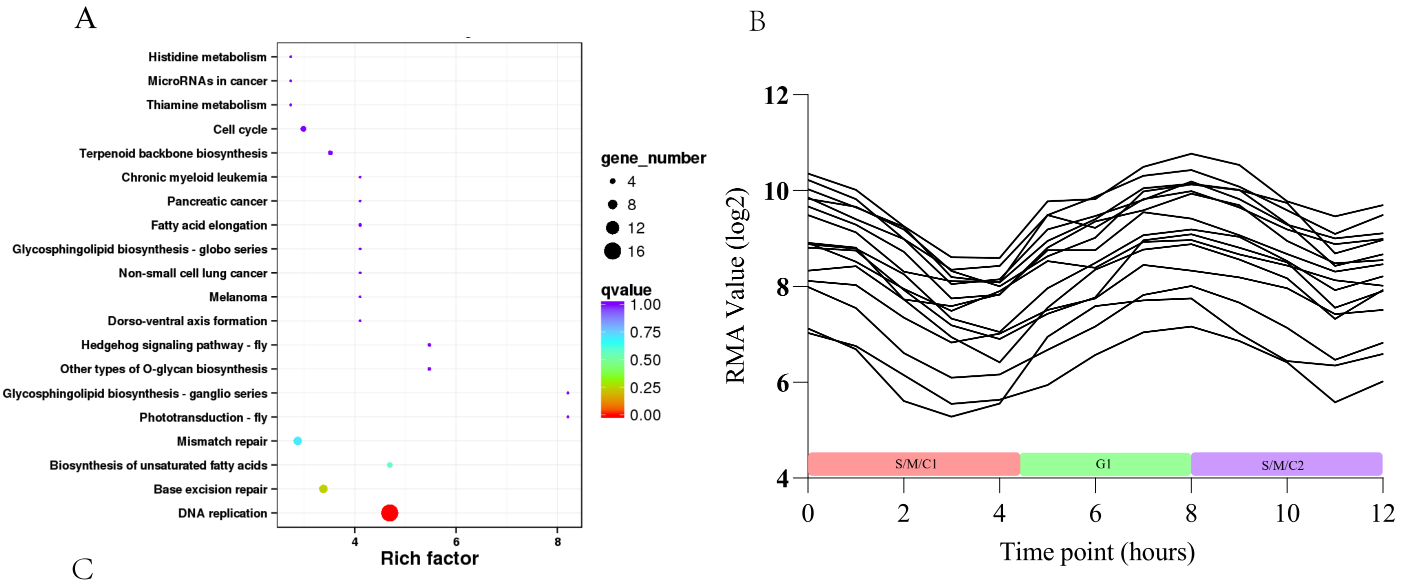


C

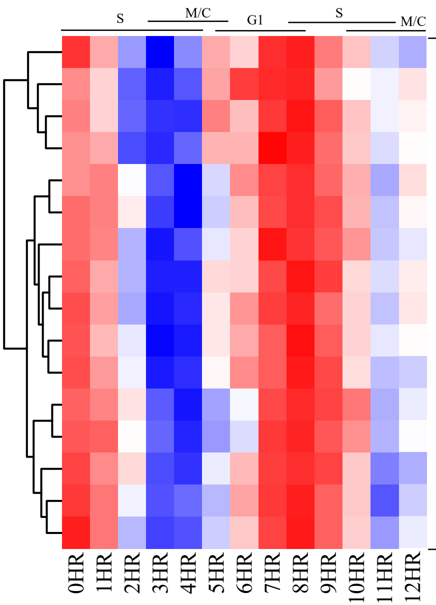


| ID | Annotation | Log2FC | Location |
|---|---|---|---|
| TGME49_208580 | DNA ligase 1 | -1.21379 | N/A |
| TGME49_220450 | ribonuclease HI protein | -1.16488 | nucleus - chromatin |
| TGME49_219700 | DNA replication licensing factor MCM4 | -2.34775 | cytosol |
| TGME49_268600 | DNA polymerase epsilon subunit B protein | -1.47627 | N/A |
| TGME49_237110 | replication factor C subunit 2 | -1.06145 | nucleus - chromatin |
| TGME49_260680 | small subunit DNA primase | -2.04396 | N/A |
| TGME49_281510 | ribonuclease H1 large subunit | -1.85709 | N/A |
| TGME49_214970 | DNA replication licensing factor | -1.54214 | nucleus - chromatin |
| TGME49_243920 | DNA replication licensing factor MCM5 | -3.52156 | nucleus - non-chromatin |
| TGME49_219860 | replication licensing factor | -1.8444 | cytosol |
| TGME49_320110 | PCNA2 | -2.04464 | cytosol |
| TGME49_203170 | OB-fold nucleic acid binding domain-containing protein | -2.03394 | N/A |
| TGME49_319860 | DNA polymerase family B protein | -2.33725 | N/A |
| TGME49_237220 | DNA replication licensing factor Mcm7 | -2.12263 | cytosol |
| TGME49_258030 | DNA polymerase | -1.34522 | nucleus - chromatin |
| TGME49_280690 | DNA polymerase epsilon subunit B protein | -2.2395 | N/A |
